# Mechanistic modeling predicts efficacy of CISH knockout in tumor-infiltrating lymphocytes with synergistic gene editing

**DOI:** 10.1101/2025.09.05.674558

**Authors:** Nikolaos Memmos, Kamran Kaveh, Beau R. Webber, Branden S. Moriarity, David J. Odde

## Abstract

Tumor-infiltrating lymphocyte (TIL) therapy is a type of adoptive cell therapy, where the lymphocytes of a cancer patient’s tumor are harvested, expanded *in vitro* using IL-2 stimulation, and then infused back into the patient[1], [2]. However, even with the use of TIL therapy, cancer cells can survive for various reasons, such as poor lymphocyte infiltration into tumors, chronic activation of the T cell receptor and the immunosuppressive tumor microenvironment[3]. Cytokine-inducible SH2-containing (CISH) protein is a negative regulator of T cell activation, and in a recent clinical trial was knocked out in TILs to improve TIL therapy efficacy[4]. A mechanistic signaling pathway model was developed to theoretically evaluate the efficacy of *CISH* knockout (*CISH* KO) in T cell activation and examine potential alternative target genes that can theoretically be targeted using multiplex gene-editing or drugs to further improve T cell activation and function[5]. Based on the results, *CISH* knockout increases the transcription of activation biomarkers IL-2 and TNF-α, but also inhibitory biomarkers such as PD1 and FasL. Using global sensitivity analysis, we also found that *GSK3B*, which is responsible for the deactivation of NFAT, is also predicted to further increase T cell activation when knocked out. In addition, it was predicted that *PDCD1*, *FAS* and *CTLA4* can be knocked out in combination with *CISH* to further enhance T cell activation and prevent exhaustion and apoptosis.

## Introduction

Adoptive cell therapy (ACT) is a form of personalized cancer treatment that consists of harvesting immune cells, expanding them *in vitro* and transferring them back to the patient to eliminate tumor cells[1]. There are two main approaches: one where immune cells are harvested directly from cancer patients to treat their own disease, i.e., autologous cell therapy, or another where immune cells are obtained from healthy donors, i.e., allogeneic cell therapy[6].

A major advancement in the efficacy of ACT was in 1976 when it was discovered that the administration of interleukin-2 (IL-2) can expand T cells *ex vivo*[2]. Moreover, Rosenberg *et al.* showed that high doses of IL-2 result in the generation of lymphokine-activated killer (LAK) cells that mediate regression of tumors in mice[7]. Donohue *et al.* examined the effect of administration of IL-2 after adoptive cell transfer[8]. Despite their initial promising results in a clinical trial, where 11 of 25 patients with different types of metastatic cancer, including melanoma, colorectal, and renal-cell, showed tumor regression after the administration of LAK, a subsequent clinical trial did not demonstrate a statistically significant benefit compared to the administration of IL-2 alone in patients with renal cell cancer[9], [10].

Tumor-infiltrating lymphocyte (TIL) therapy is a novel cancer immunotherapy in which the patient’s own immune cells, present in the tumor microenvironment, are harvested, expanded *in vitro* and reinfused back to the patient to fight cancer. TILs after expansion with IL-2 showed 50 to 100 times higher efficacy compared to LAK cells in lung and hepatic metastatic tumors, although only when combined with pretreatment using cyclophosphamide[11]. The first example of TIL therapy was in 1988, when Rosenberg *et al.* used autologous TILs along with the administration of IL-2 and pretreatment with cyclophosphamide to treat 20 patients with metastatic melanoma[12]. This study showed that the administration of TILs resulted in a positive response in 9 out of 15 patients who were not treated with IL-2 before and in 2 of 5 patients who were treated with IL-2 in the past, compared to IL-2 and cyclophosphamide alone. A significant milestone in TIL therapy happened in 2002 when it was shown that the use of nonmyeloablative chemotherapy regimen as conditioning to achieve lymphodepletion before TIL infusion in patients with metastatic melanoma causes tumor regression[13].

Neoantigen-reactive TIL therapy leverages the subset of tumor-infiltrating lymphocytes that are specific to the neoantigens expressed by tumor cells. Selection of neoantigen-reactive TIL pools can be accomplished using immunoassays to evaluate reactivity against candidate neoantigens nominated by next-generation sequencing of patient tumors[14]. Neoantigen-reactive TIL therapy has shown the potential to cure metastatic cancer. Zacharakis *et al.* presented the case of a patient with metastatic breast cancer who was treated with TILs specific to neoantigens created by nonsynonymous mutations in four genes, *SLC3A2, KIAA0368, CADPS2* and *CTSB*, and led to complete regression that persisted for more than 22 months[15]. Tran *et al.* published a case report of a patient with metastatic colorectal cancer. The patient was treated with T cells that target a neoantigen created by KRAS G12D[16]. All seven metastatic sites in the lungs regressed but one of them relapsed after 9 months of therapy because of the loss of chromosome 6 that encoded HLA-C*08:02 protein, which is necessary for the T cells to recognize the KRAS G12D tumor antigen. In 2024, the FDA approved the first TIL therapy for patients with melanoma who have been treated before with PD1 inhibitor[17].

Despite the success of neoantigen-reactive TIL therapy, it is not yet the norm. Zacharakis *et al.* examined the efficacy of the therapy in conjunction with pembrolizumab in patients with metastatic breast cancer[18]. TILs were isolated from 42 patients but only 6 of them were found eligible to be treated. From these patients, only 1 had a complete response which persisted over 5.5 years, 2 partial responses at 6 and 10 months, and tumor regression in 3 patients. This may be attributed to the immunosuppressive properties of the tumor microenvironment in solid tumors, for example the poor infiltration of lymphocytes, the expression checkpoint inhibitors, poor neoantigen presentation and chronic T cell receptor (TCR) activation[3]. To counteract these effects, there have been intensive efforts to override the intrinsic checkpoints that limit T cell activation pathways.

Cytokine-inducible SH2-containing (CISH) protein is a member of suppressors of cytokine signaling (SOCS) family and a novel intracellular checkpoint[19]. It has been shown that all eight members of the SOCS family, which are SOCS1-7 and CISH, play a key role in immune regulation[20], [21], [22], [23]. Delconte *et al.* observed that CISH protein is a negative regulator of IL-15 in natural killer cells, and its deletion increases sensitivity to IL-15[24]. The deletion of *CISH* also increased production of IFN-γ and cytotoxicity against tumors, because of its interaction with JAK1 protein in the JAK-STAT signaling pathway[22], [23], [25], [26], [27]. Palmer *et al.* also demonstrated that CISH is responsible for the ubiquitination and degradation of PLC-γ1 in T cells after TCR activation[19]. *In vivo* experiments showed that knocking out CISH protein increased the activation of T cells by enhancing the translocation of NFAT and NFκB in the nucleus, while it also increased the phosphorylation of ERK which contributes to higher T cell activity[28]. These results led to the development of genetically engineered TILs using CRISPR/Cas9 *CISH* knockout, resulting in a human phase I/II clinical trial (NCT04426669) to treat metastatic gastrointestinal cancer[4].

Here, we present a modeling study where we examine whether *CISH* KO can theoretically increase the activation and function of T cells, and whether there are other genetic modifications that could be used instead of, or in combination with *CISH* KO to achieve a significant increase in T cell activation. A mechanistic model has been previously developed using ordinary differential equations (ODEs) that captures the effects of NFAT with respect to important biomarkers for T cell activation, such as IL-2, TNF-α, CTLA-4 and Fas ligand (FasL), as described in Shin *et al.*[5]. Based on their model, we expanded it to include PD1, CISH, and PLCγ1 proteins, as well as more protein interactions, to capture the effect of *CISH* KO in T cell function. IL-2 and TNF-α are both responsible for the activation and expansion of T cells[29], [30], [31], [32]. In addition, TNF-α plays a major role in recruiting other immune cells in the tumor site[33], [34]. On the other hand, PD1 acts as an immune checkpoint to prevent the overactivation of T cells and contributes to T cell exhaustion, while inhibiting the activation of TCR, PI3K and Ras signaling[35], [36], [37]. Furthermore, CTLA-4 has an inhibitory action on T cells by competing with CD28 for binding to the B7 protein on antigen presenting cells (APCs) and it inhibits the action of PP2A[38], [39], [40]. The role of FasL is to maintain immune homeostasis by inducing activation-induced cell death (AICD) in T cells[41].

Our results predict that *CISH* KO will be highly effective in elevating the expression of both activation and inhibitory markers. Moreover, our results indicate that multiplex gene-editing of *CISH*, *PDCD1*, *FAS*, *CTLA4* may improve the efficacy of TILs, and also suggest that inhibition of GSK3β protein, which increases proliferation of T cells, could further improve T cell function in this context[42], [43], [44].

The paper is organized as follows: We first describe the model, including the kinetic framework. We then perform parameter estimation using experimental datasets from the literature to establish a calibrated baseline model. Next, we conduct global sensitivity analysis to identify the most influential parts of the pathway that control the expression of IL-2, TNF-α, PD-1, CTLA-4, and FasL proteins, which we consider key outputs of the system. Using these findings, we perform *in silico* experiments to evaluate the effect of *CISH* knockout alone and in combination with *PDCD1* and *GSK3B* knockouts across heterogeneous virtual T cell populations to assess potential synergistic therapeutic strategies. Finally, we discuss the biological and clinical implications of multiplex gene editing for TIL therapy, as well as model limitations and future directions.

## Model

The formulation of the CISH model developed here is based on the paper of Shin *et al.*[5], originally developed to study activation-induced cell death (AICD) in T cells. Since our model is also centered around the activation and effect of NFAT, we reasoned that Shin *et al*. was a reasonable choice for a starting model framework. In our study, the Shin et al. model was modified to integrate PLCγ1, CISH and PD1 and their interactions with other proteins **(Figure 1**). Enzyme-catalyzed reactions and indirect cascade effects were modeled using Michaelis-Menten reaction kinetics. For example, calcineurin activation, which is triggered by calcium release in the cytosol through the endoplasmic reticulum after the phosphorylation of PLCγ1, was simplified using a Michaelis-Menten equation (**Supplementary Table A.1,** reaction 6) to reduce model complexity. Similarly, the activation of TAK1, which is induced by TNF-α via tumor necrosis factor receptor (TNFR) was reduced to a single Michaelis-Menten equation (**Supplementary Table A.1**, reaction 43).

**Figure 1.**
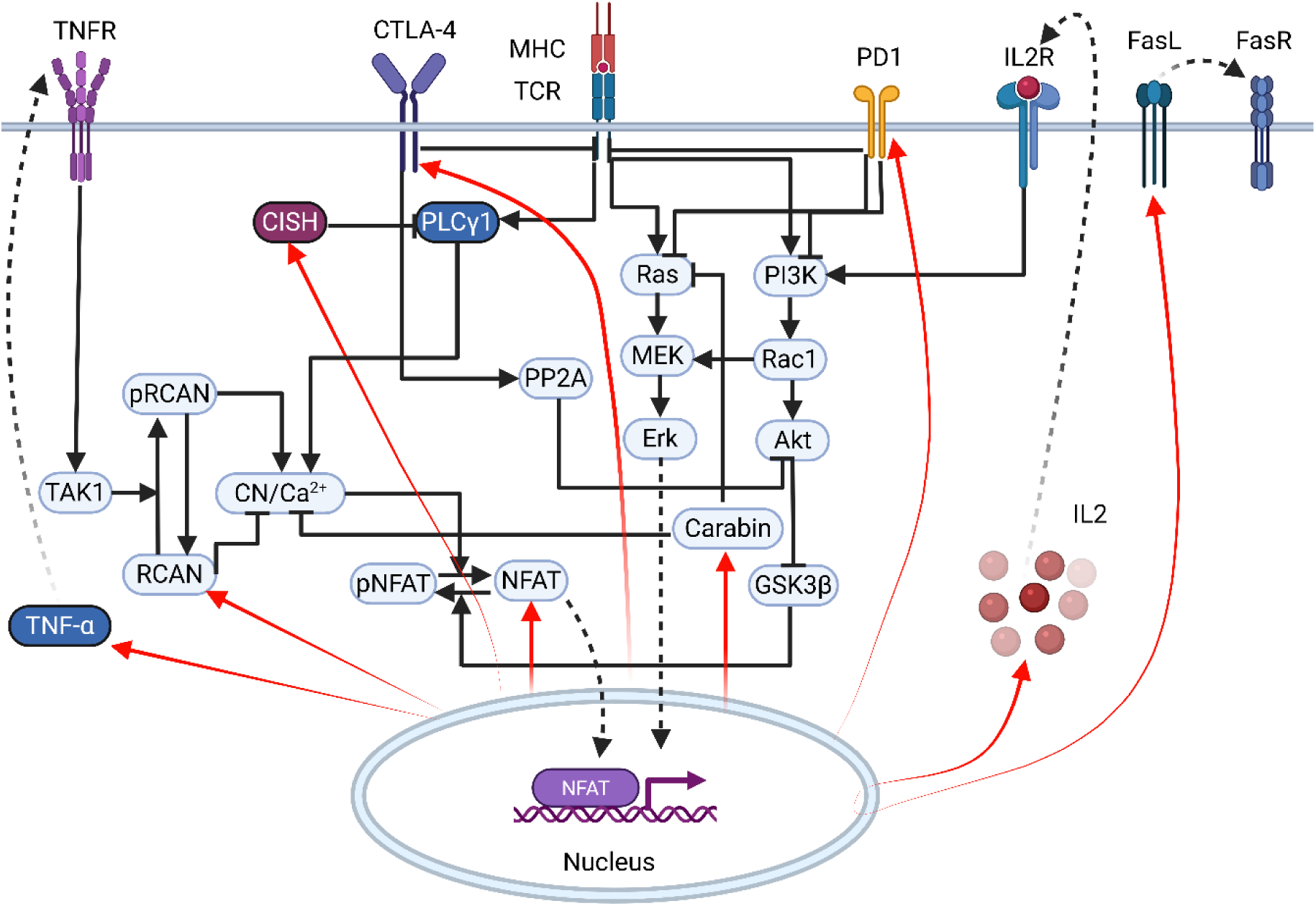
Model schematic for NFAT activation and downstream effects upon TCR activation. Black solid arrows and T-arrows indicate the activation and inhibition of proteins, respectively. Red arrows represent the expression of proteins that are dependent on NFAT transcription factor acting in the nucleus. Dashed black arrows indicate the translocation of proteins to the nucleus or in the extracellular space.

Moreover, the Michaelis-Menten equations were decomposed into a two-step reaction, comprising a reversible binding step (𝐸 + 𝑆 ↔ 𝐸𝑆) and an irreversible catalytic step (𝐸𝑆 → 𝐸 + 𝑃), to enhance model stability during sensitivity analysis and maintain mass conservation. mRNA production rates were modeled as zeroth-order reactions (i.e., occurred at a constant rate), while degradation was modeled as a first-order reaction, with rates proportional to the concentration of the species being degraded. To account for the role of NFAT in protein transcription, Hill functions were used to describe the target gene mRNA expression dependence on NFAT. Mass-action kinetics were used to model association and dissociation events (**Supplementary Table A.1**).

The strength of the model of Shin *et al.* lies in its detailed representation of the feedback loops that are involved in the NFAT expression[5]. In addition, our model incorporates the NFAT-mediated feedback loop for CISH expression, wherein NFAT activation enhances CISH transcription, and subsequently it leads to the ubiquitination of PLCγ1. Additionally, the model includes the regulation of PD1 expression via the NFAT translocation to the nucleus and the inhibitory effect of PD1 protein to the TCR, Ras and PI3K signaling. Our model consists of 160 free parameters and 74 ODEs and uses the same initial conditions as the original Shin *et al*. model. For the rest of the species, they were assumed to be 10 nM for the free intracellular proteins, 0 nM for the protein complexes, 0.1 nM for the surface proteins (e.g. CTLA-4) and mRNAs. The model was implemented in the Julia programming language[45].

## Methods

### ODE solvers

The DifferentialEquations.jl package in Julia was utilized in this study, since it provides a wide range of ODE solvers[46]. To solve the ODE system, the QNDF or the Rosenbrock23 methods were used[46], [47], [48]. The Rosenbrock23 solver was paired with the Tsit5 solver using an auto-switching algorithm to enable the switching between these two solvers, based on the stiffness of the ODE system[49]. The QNDF solver was used during the parameter estimation step, while the Rosenbrock23/Tsit5 solvers were used during the global sensitivity analysis and steady-state simulations because of their ability to take larger time steps; thus reducing the computational cost[50]. To further improve computational efficiency during global sensitivity analysis and steady-state simulations, numerical differentiation with central differencing was implemented. For parameter estimation, automatic differentiation was employed, providing higher accuracy but at a greater computational cost[51].

### Parameter estimation

Published *in vitro* experimental data were used to establish baseline parameter values [5], [39], [52], [53], [54], [55]. The lower and upper bounds of the parameter ranges are given in **Supplementary Table A.3**, while the estimated parameter values in **Supplementary Table A.4**. Because of large number of model parameters and the limited experimental data, many of the model parameters are non-identifiable[56]. To tackle this issue, Fides package for interior trust region reflective for boundary constrained optimization was used[57]. The objective function of the optimization problem is given by

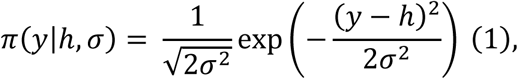

where 𝑦 is the experimental measurement, ℎ is the model observable and 𝜎 the standard deviation of the noise. A common problem in intracellular networks is the mismatch between the model outputs and the form of the experimental measurements, which in many cases are normalized or have different units. For this reason, a linear mapping between the observables and the experimental measurements was used[58], given by

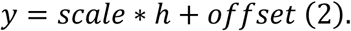

To estimate the gradient of the objective function with respect to the model parameters, forward automatic differentiation method was used. For computing the Hessian matrix, the Gauss-Newton method was used[59]. PEtab.jl library in the Julia programming language was used to implement parameter estimation[60]. The absolute and relative tolerances of the ODE solver were set to 10^-3^ and 10^-6^ respectively.

### Global Sensitivity Analysis

To identify the sensitive parameters, the Morris method was used[61]. The Morris method is based on the calculation of elementary effects (𝐸𝐸), which are defined as

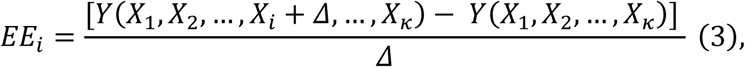

where 𝑌 is the output of the model, 𝑋_𝑖_, 𝑖 = 1, … , 𝑘 are the parameters of the model and 𝛥 is the step size, which takes values in 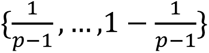 and 𝑝 is the number of grid points in the parameter hyperspace. To sample the parameter hyperspace, 𝑟 trajectories of 𝑘 + 1 points are created, with 𝑘 elementary effects per trajectory, one for each parameter, and total 𝑟(𝑘 + 1) sample points. The trajectories are selected from a superset of trajectories to have the biggest spread in the parameter space[62]. In our analysis 20,000 trajectories, for each parameter, were selected from a set of 200,000 trajectories.

The Morris method estimates the following sensitivity measures; 𝜇, 𝜇^∗^ and 𝜎,

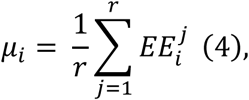

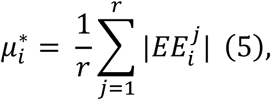

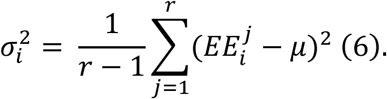

Both 𝜇 and 𝜇^∗^are the means of the elementary effects and the absolute values of them respectively, and they are used to assess the overall influence of a parameter, while 𝜎^2^is the variance of the distribution of the elementary effects and measures the non-linear and interactions with other parameters. Compared to 𝜇, the use of 𝜇^∗^prevents potentially cancelling effect between positive and negative effects in the cases of non-monotonicity or interactions with other parameters.

## Results

### Model calibration

The model was calibrated using experimental time series data from multiple sources for CISH, PLCγ1, phosphorylated PLCγ1 (pPLCγ1), Akt1, phosphorylated MEK (pMEK), phosphorylated ERK (pERK), TNF-α, IL-2, mRNA coding for IL-2 (mIL-2), NFAT, PD1, CTLA-4, and phosphorylated TCR (pTCR)[5], [39], [52], [53], [54], [55]. In all instances, wild-type T cells were stimulated with anti-CD28 and anti-CD3 antibodies. The CISH, PLCγ1 and phosphorylated PLCγ1 data were extracted from western blots using Fiji Image software, while the remaining data were extracted using WebPlotDigitizer[63], [64]. The initial stimulation of the T cell was modeled as a rectangular pulse function with a 2-minute duration. Despite the large number of free parameters and the limited experimental data, Fides algorithm performed well in fitting the model to the experimental data. To estimate the parameters, 10000 starts from different initial points in the parameter space were used (**Supplementary Figure A.1**).

NFAT, IL-2, and mIL-2 displayed longer peaks in protein expression following TCR stimulation (**Figures 2A-C**), compared to pAkt, pMEK, pERK, and pPLCγ1, which all exhibited a rapid increase followed by a fast reduction in intracellular concentrations (**Figures 2D-F**, **Figure 2H**). In contrast, CISH and TNF-α concentrations demonstrated a consistent increase over time (**Figure 2I, 2K**). However, it is important to note that the datasets for CISH, PLCγ1, and pPLCγ1 contained only four data points at early time points, potentially limiting the ability of the model to make accurate predictions at later times. PD1 concentration displays a much slower decay rate compared to the rest (**Figure 2L**). Additionally, CTLA-4 exhibits an initial peak after TCR activation and then a decrease that leads to a persistent expression (**Figure 2J**).

**Figure 2.**
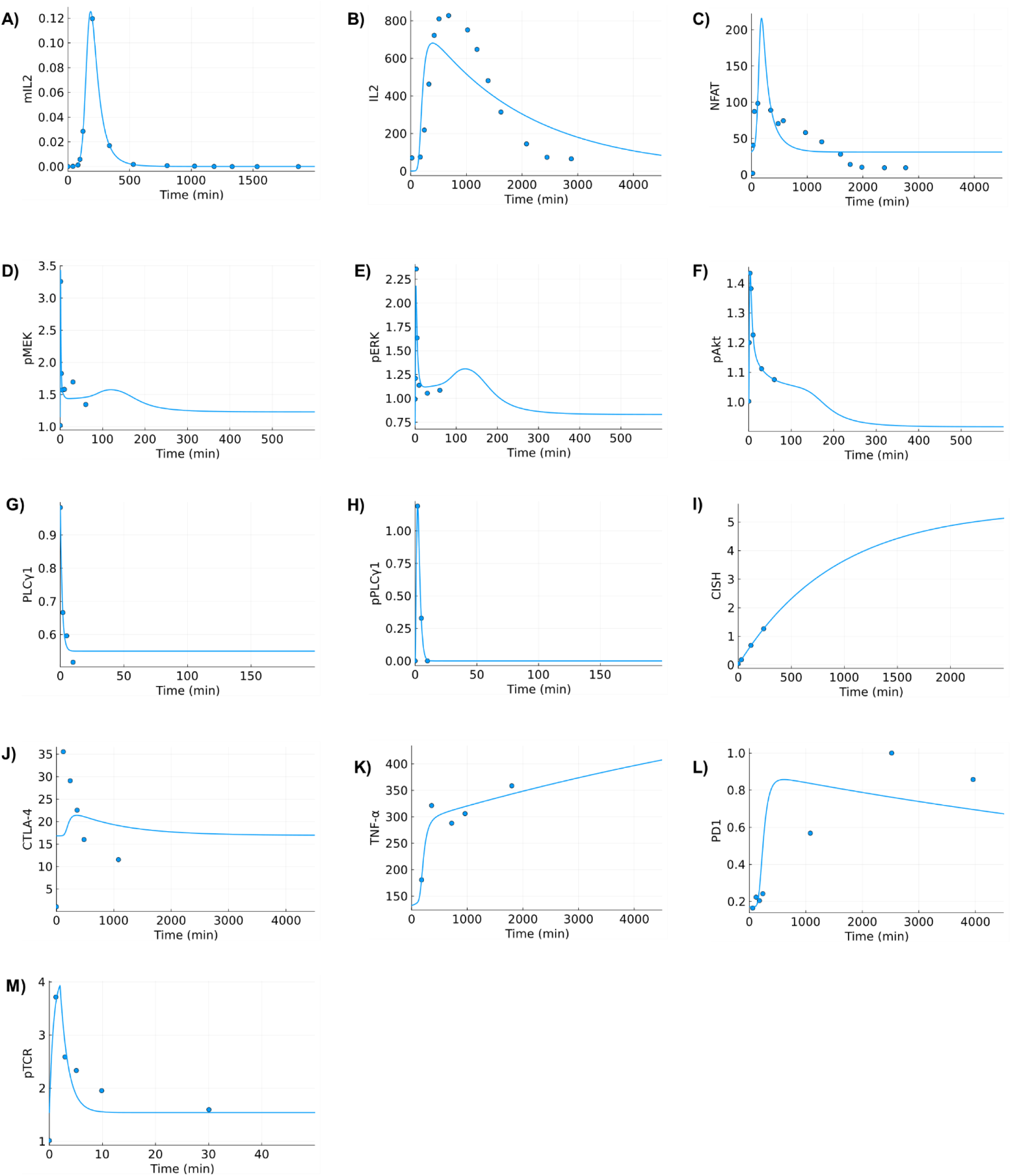
Model calibration based on **A)** mRNA IL2 (mL2), **B)** IL2, **C)** NFAT, **D)** pMEK, **E)** pERK, **F)** pAkt, **G)** PLCγ1, **H)** pPLCγ1, **I)** CISH, **J)** CTLA-4, **K)** TNF-α, **L)** PD1, **M)** pTCR. The blue dots correspond to the experimental data obtained from references [5], [39], [52], [53], [54], [55] and the solid lines correspond to the model output. Equation (2) was used for the mapping between the model output and the experimental data.

The large number of model parameters and limited experimental data hinder the predictive power of the model due to identifiability and overfitting issues. In our case, parameter fitting is crucial for establishing baseline values for the intracellular biological processes that we consider therapeutically unmodifiable, primarily the baseline generation and degradation rates of proteins and mRNAs, *kgen_i_* and *kdeg_i_* , respectively (**Supplementary Table A.1**), which were held constant for the rest of the analysis. In the case of proteins that depend on the NFAT upregulation, CISH, RCAN, Carabin, CTLA-4 and PD1, and the expression of mRNA coding for NFAT, both the production rate of mRNAs and the proteins were considered as free parameters. Moreover, the Michaelis-Menten parameter, *km_i_*, (**Supplementary Table A.1**) was kept constant based on the estimated values from the parameter fitting step. Using this approach, the number of free parameters was reduced from 160 to 71, which further reduced the computational cost of the sensitivity analysis. Furthermore, by limiting the number of free parameters during the rest of the analysis, we were able to obtain a better understanding of the important parts of the pathway.

### Critical parts of the pathway

To identify if *CISH* is a good candidate for targeting and what other parts of the pathway can potentially be targeted independently or in conjunction with *CISH* KO, global sensitivity analysis (GSA) was performed using the Morris method. The outputs of Morris method are the 𝜇_i_^∗^(**Equation 5**), which measures the influence of each parameter, and 𝜎_i_^2^(**Equation 6**) that corresponds to the variance of the output with respect to parameter 𝑖, which corresponds to a nonlinear or interacting behavior of the parameter.

As an output for the sensitivity analysis, the steady-state concentrations of IL-2, TNF-α, PD1, FasL and CTLA-4 were used. It was assumed that T cells are, on average, in a continuously activated state in the tumor microenvironment[65], [66]. Compared to other global sensitivity analysis methods, for example Sobol, the Morris method is less accurate in terms of the ranking of the parameters, but because of the prohibitively high computational cost of the Sobol method due to the number of free parameters and steady-state simulations, the Morris method was used for the purpose of this study.

The expression level of IL-2 protein exhibits high sensitivity to the production velocity of CISH (*v33*) and the deactivation of calcineurin, *v6a*, with values of 𝜇_𝑣33_^∗^ and 𝜇_𝑣6𝑎_^∗^ equal to 3.36 and 3.14 respectively (**Figure 3A**). Furthermore, the high variances of v33 and v6a parameters indicate a possible interaction with other parameters. The production rate of PD1, *v35*, and the dephosphorylation rate of pGSK3β, *v27*, have a lower effect on IL-2 expression, but parameter v35 shows a high variance (𝜎_v35_^2^ = 142.9), similar to the production velocity of mNFAT, *v6*, which corresponds to the positive self-feedback loop of NFAT regulation and has a high variance, with a value 186.29. A summary of the parameters and their meaning is shown in **Table 3.1**.

**Figure 3.**
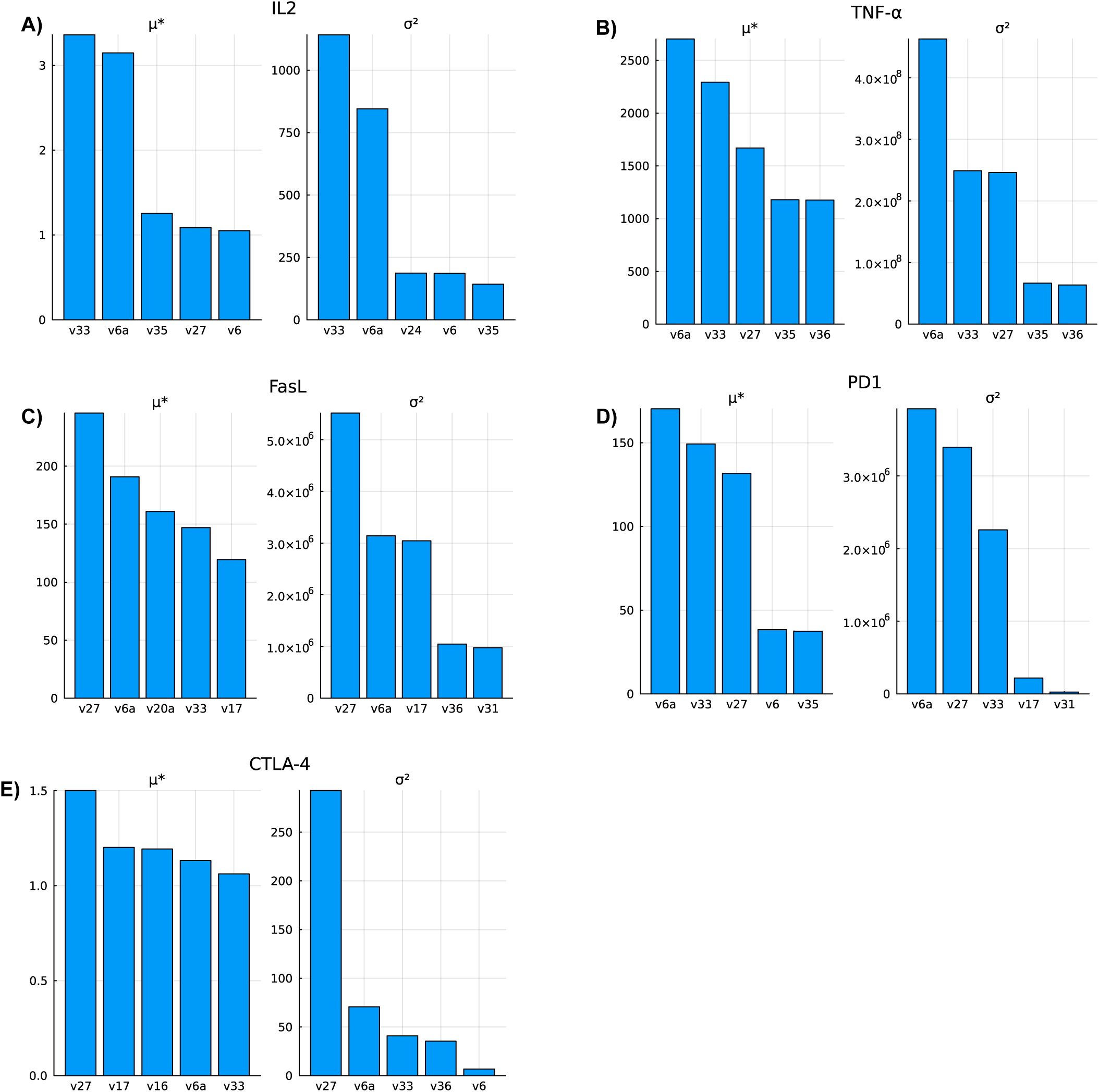
Parameter sensitivity analysis using the Morris method for **A)** IL-2, **B)** TNF-α, **C)** FasL, **D)** PD1 and **E)** CTLA-4 steady state concentrations under constant activation of the TCR. The mean, 𝜇_i_^∗^, is used to quantify the sensitivity of each output with respect to variations of a parameter, and the variance, 𝜎_i_^2^, for the nonlinear behavior or interactions with other parameters.

**Table 1.** Sensitive parameters based on Morris global sensitivity analysis method and their biological meaning.

| Parameter | Meaning |
| --- | --- |
| v33 | Production of CISH protein |
| v6a | Deactivation of calcineurin |
| v35 | Production of mPD1 |
| v36 | Production of PD1 |
| v27 | Dephosphorylation of pGSK3β |
| v6 | Production of mNFAT |
| v24 | Dephosphorylation of ERK |
| v20a | Production of FasL |
| v16 | Production of mCTLA-4 |
| v17 | Production of CTLA-4 |
| v31 | Phosphorylation of inactive IL2R |

TNF-α is extremely sensitive to the calcineurin deactivation, CISH production and dephosphorylation pGSK3β due to the high values of 𝜇_i_^∗^ (**Figure 3B**), while the parameters also exhibit variances on the order of ∼10^8^. Furthermore, the parameters *v27*, *v6a* and *v33* appear to influence FasL, PD1 and CTLA-4 (**Figures 3C-E**). According to the sensitivity analysis, PD1 and FasL are susceptible to changes in CISH production, dephosphorylation rate of pGSK3β, and deactivation of calcineurin, but also these parameters probably interact with other parameters. On the other hand, CTLA-4 appears to be less sensitive to these parameters. In the case of CTLA-4, the dephosphorylation rate of *v27* shows a high variance, which stems from interactions with other parameters or non-linear effects. Parameter *v17*, which influences FasL and CTLA-4, corresponds to the production of CTLA-4. An important observation is parameter *v31*, which corresponds to phosphorylation of IL-2 receptor (IL2R), has a high variance in FasL and PD1 respectively (variances equal to 9.8*10^5^ and 2.6*10^4^), suggesting that the IL2R-PI3K-Rac1-Akt axis is potentially important.

Based on the global sensitivity analysis we confirm that theoretically *CISH* is an excellent target for genetic KO, via gene editing, as it is implicated in the expression of multiple distinct markers: *CISH* KO can increase the transcription of IL-2, TNF-α, PD1 and FasL. CTLA-4 is the most robust to parameter changes, compared to the other markers. Moreover, calcineurin appears to have an even more important role, since it directly acts on NFAT activation, while *PDCD1* and *GSΚ3Β* gene-editing can also be used in conjunction with *CISH* knockout.

### Efficacy of CISH KO

To analyze the long-term effects of *CISH* KO in T cells, 10000 cells were generated, under constant activation and with different parameter values, using sampling from a log-normal distribution with a mode equal to the fitted value of each parameter,

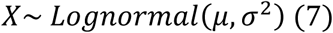

where 𝜇 and 𝜎 are the mean and the standard deviation of ln(𝑋) ∼𝑁(𝜇, 𝜎^2^) distribution. The mode of the lognormal distribution is given by:

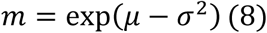

It was assumed that 𝜎 is equal to 0.3; thus all the sampled parameters will have the same coefficient of variation (CV=0.307), which is calculated based on the following equation[67]:

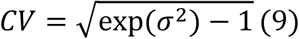

Steady-state levels of IL-2, TNF-α, PD-1, FasL and CTLA-4 were used to evaluate the long-term activation and exhaustion, as persistent antigen exposure changes the expression of these markers in tumor-resident TILs. From the simulations, it was observed that there is a large variability in the wild type T cells, with FasL exhibiting high expression in some cases and low expression in others. After knocking-out *CISH*, TNF-α and IL-2 increase, but also PD1, CTLA-4 and FasL, which is attributed to the rise of NFAT (**Figure 4**). This simultaneous rise in all markers can potentially explain the inability of *CISH* KO by itself to lead to tumor regression in mouse models of B16 melanoma[4], [28].

**Figure 4.**
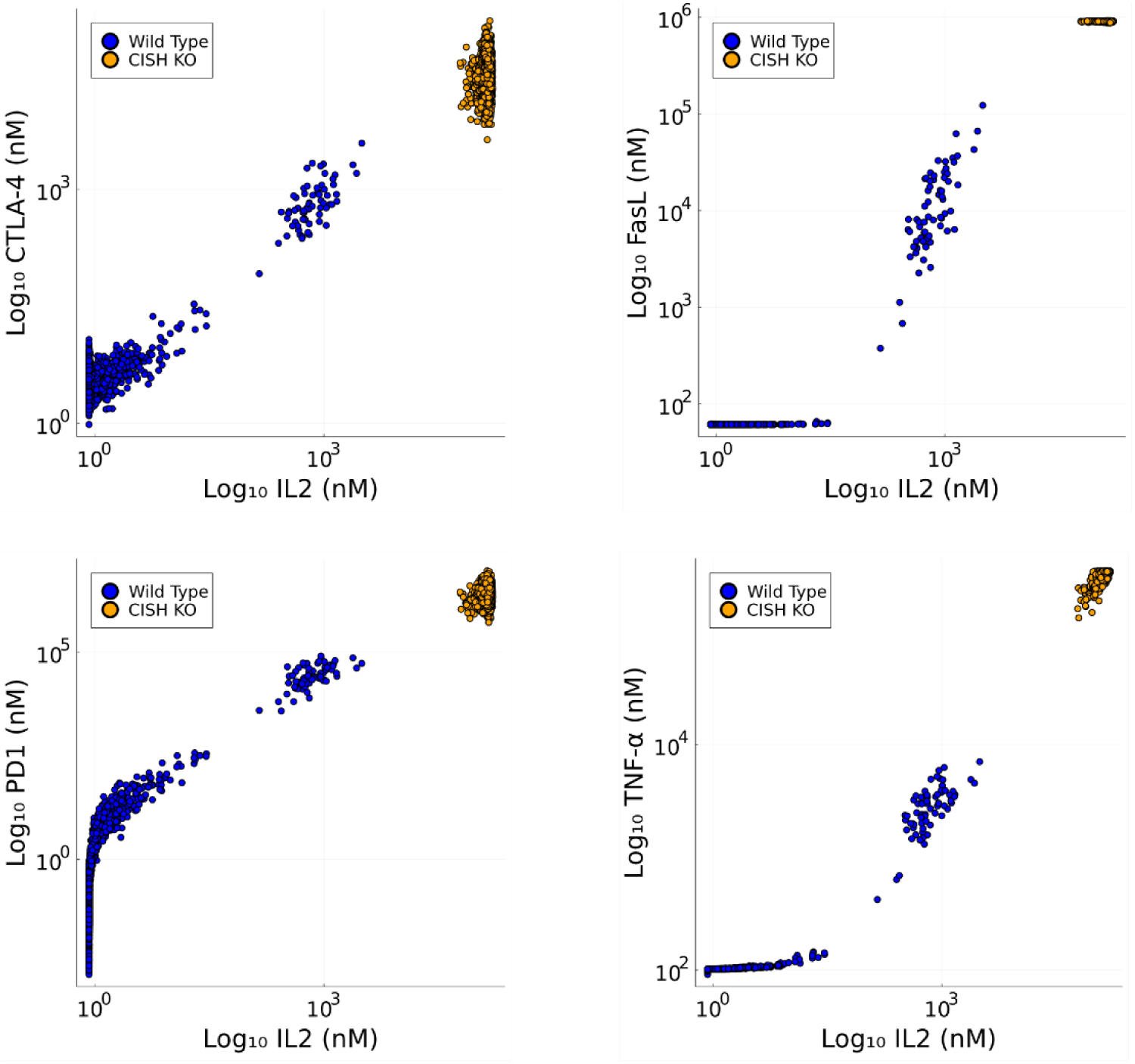
*In silico* steady state simulations for wild type and CISH KO T cells. CISH KO increases the expression of all markers. CISH KO simultaneously increases the expression of IL-2, TNF-α, PD1, FasL and CTLA-4.

### Therapy recommendations

As noted in Palmer *et al.*, *CISH* KO by itself was not sufficient to reduce tumor size in B16-melanoma bearing mice, but the combination of *CISH* KO and anti-PD1 block therapy had a high anti-tumoral efficacy[28]. Furthermore, Arthofer *et. al.* reported an increase in tumor cell killing after inactivation of both *CISH* and *PDCD1*[68]. In addition, the global sensitivity analysis of the model suggests the dephosphorylation rate of pGSK3β to be important in some of these proteins. GSK3β is pivotal for deactivating NFAT via phosphorylation, since NFAT is inactivated when phosphorylated.

To test this in the model, the production rates of CISH and PD1 mRNAs, *v32* and *v35* respectively, were set equal to zero to simulate simultaneous knockout of *CISH* and *PDCD1* via multiplex gene editing[69]. In the case of simulated *GSK3B* knockout, both the initial conditions of GSK3β and pGSK3β were set to zero. Similarly to the previous section, 10000 cells for each case were simulated in a steady state. The synergistic inactivation of *CISH* and *PDCD1* appears to leads to a higher expression of IL-2, compared to *CISH* KO alone, while *PDCD1* KO did not offer a significant improvement, as the IL-2 concentration distribution in this case is similar to the wild type cells (**Figure 5A**), as was also shown by Cano *et al.*[70]. When GSK3β is knocked-out in combination with *CISH* and *PDCD1*, the production of IL-2 is further increased in the model as evidenced by the higher IL-2 concentrations in the distribution in this case. In addition to the increase of IL-2, the multiplex gene knockout of *CISH*, *PDCD1* and *GSK3B* is predicted to result in a better outcome for TNF-α, in line with the observation from the sensitivity analysis that TNF-α is the most sensitive to these changes (**Figure 5B**). However, FasL increases significantly in *CISH* KO and the combinations of *CISH* KO + *PDCD1* KO or *CISH* KO + *PDCD1* KO + *GSK3B* KO have a modest effect (**Figure 5C**). The simulations suggest that CTLA-4 increases significantly with the combination of *CISH* KO with *PDCD1* KO and *GSK3Β* KO (**Figure 5D**).

**Figure 5.**
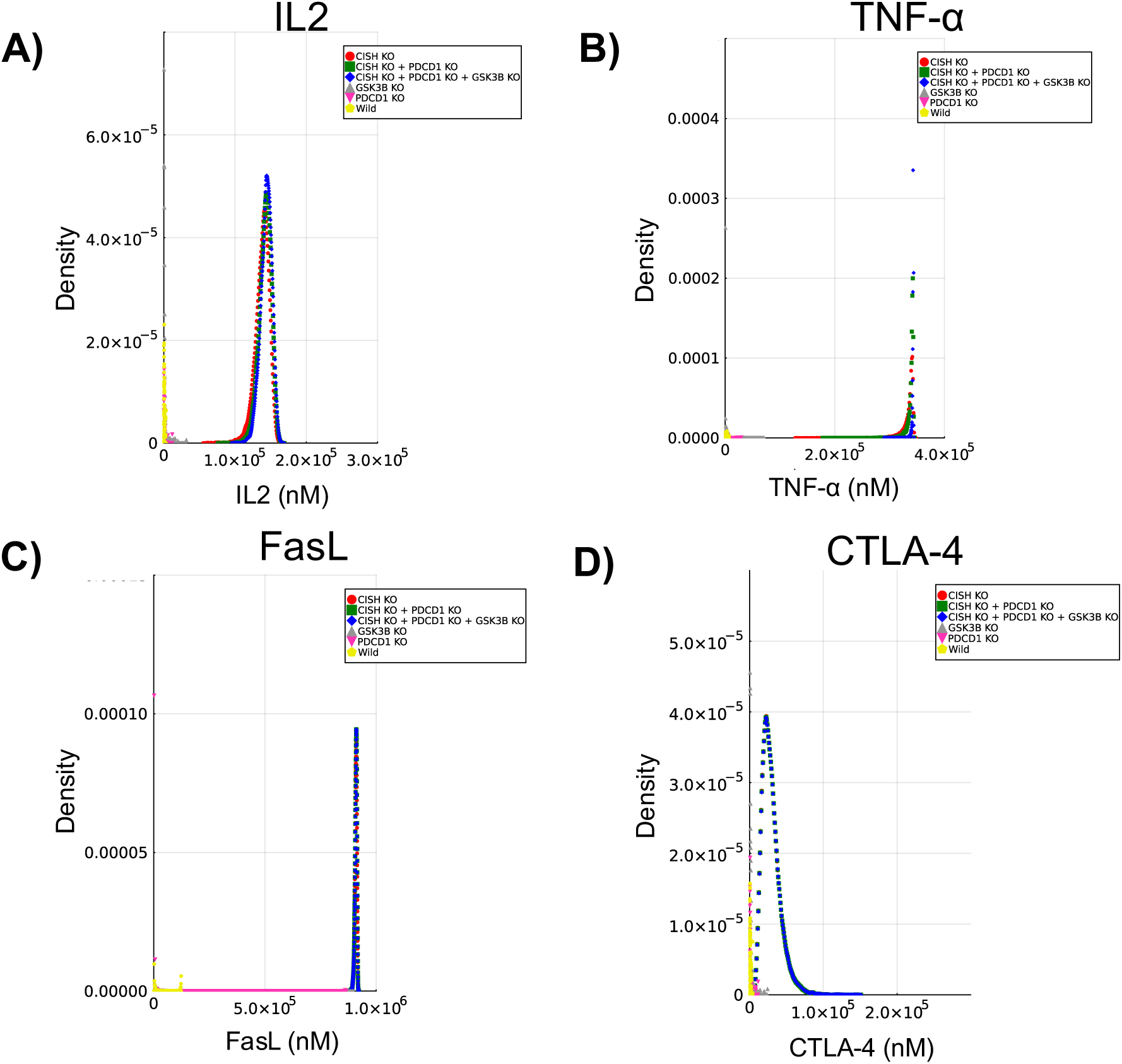
Steady-state concentration distributions of **A)** IL-2, **B)** TNF-α, **C)** FasL and **D)** CTLA-4 for different therapy combinations. CISH KO + PD1 KO + GSK3β KO leads to an increase in IL-2, TNF-α and FasL. CTLA-4 only requires CISH knockout to have higher expression, whereas the other outputs are further increased at steady-state by combining CISH KO with one or more other gene knockouts, such as PD1 and/or GSK3β.

According to these results, *CISH* KO is predicted to lead to the biggest increase in the biomarkers, compared to the individual gene knockouts, since it acts directly on PLC-γ1 and it has been shown that PLC-γ1 deficiency in T cell impairs their activation upon TCR stimulation[71]. Our model-based finding that CISH is an especially important negative regulator/checkpoint in T cell activation aligns with recent experimental studies that similarly point to an especially important role for CISH[70]. CISH KO results in an increase in the transcription of IL-2 and TNF-α, with the latter being the most sensitive in this change. Additional inactivation of PDCD1 and GSK3B may further enhance T cell activity, although these predictions should be interpreted as hypothesis-generating rather than definitive therapeutic strategies. Because of the increase of FasL expression, which can lead to AICD when it binds to Fas receptor (FasR), our simulations suggest that, in theory, a broader multiplex strategy targeting *CISH*, *PDCD1*, *GSK3B*, *CTLA4* and *FAS* might boost T cell reactivity, prevent exhaustion from PD1 and CTLA-4 and AICD from FasL. The benefits of targeting CTLA-4 and PD1 have been shown in numerous studies, including clinical trials[72], [73], [74], [75]. On the other hand, GSK3β is less popular as a potential target, despite its role in the inhibition of T cell proliferation as a result of the phosphorylation of NFAT[42], [43]. Previous studies demonstrated that GSK3β inhibition led to better activation of T cells by downregulating PD1 and LAG3 proteins, while increasing the expression of transcriptional regulator TBET[44], [76], [77], [78], [79], [80], [81]. These effects appear to balance the reduced T cell motility after the inhibition of GSK3β[82].

## Discussion

Adoptive cell therapy, such as TIL therapy, is an emerging form of cancer therapy that leverages antitumoral lymphocytes to fight against cancer cells, resulting in tumor regression or even eradication[1], [7], [8]. Neoantigen TIL therapies manage to mitigate this barrier with higher cancer cell specificity, as has been reported in cases like metastatic bladder and breast cancers[83]. Despite the progress, there are many other limiting factors, such as the tumor microenvironment, which leads to suppressed proliferation and migration of T cells, low expression of neoantigens and the major histocompatibility complex (MHC) on cancer cells[3].

Traditional checkpoint inhibitors, such as anti-PD1 and anti-CTLA-4 antibodies, have shown beneficial therapeutic outcomes, but the high patient and tumor variability of PD1 and CTLA-4 expressions limit their efficacy[68], [72], [75], [84], [85], [86], [87], [88], [89]. CISH protein is an intracellular checkpoint that is responsible for the ubiquitination of PLC-γ1 and multiple studies have reported that *CISH* knockout enhances TIL reactivity against cancer cells. Previous studies have established CISH as an intracellular checkpoint limiting T cell activation, but the mechanistic basis for the variable efficacy of CISH knockout has remained unclear. Palmer *et al.* found that *CISH* knockout alone does not lead to tumor regression, because of the increase in PD1, whereas a combination of *CISH* knockout and anti-PD1 therapy resulted in tumor regression in mice[28]. In this study, we developed an ODE signaling pathway model to simulate the roles of CISH and NFAT in T cell activation. Our model was based on the model of Shin *et al.*, and modified to capture the effect of CISH in the expression of functional output biomarkers[5], including IL-2, TNF-α, PD1, CTLA-4 and FasL.

To estimate the parameters of the model, *in vitro* data from the literature have been used, where in all cases, T cells were stimulated with anti-CD28 and anti-CD3 antibodies to induce activation[5], [39], [53], [54], [55], [89]. There are significant differences in the transient behavior among the expression of proteins, for example NFAT and IL-2 exhibit a wider peak, compared to pAkt, pMEK, pERK, and pPLCγ1, which have a faster response followed by rapid decay. On the other hand, CISH and TNF-α increase over time, but in the case of CISH, we only had four data points in early times; thus we cannot be confident that this is the actual behavior at longer times post-activation. Furthermore, this limitation applies to pPLCγ1 and PLCγ1, while CTLA-4 expression exhibits an initial peak and then settles to a steady state value. Despite the limited amount of data for some of the outputs, the parameter estimation mostly served the purpose of establishing the baseline values for the protein degradation and generation rates, *kdeg_i_* and *kgen_i_* respectively, as well as the Michaelis-Menten constants, *km_i_*.

To evaluate which parts of the pathway influence the outputs of the model the most, global sensitivity analysis was performed, using the Morris method[61]. The sensitivity analysis was used to identify the most influential parameters with respect to the steady-state concentrations of IL-2, TNF-α, FasL, PD1 and CTLA-4, under constant stimulation of TCR. The estimated baseline parameters of *kdeg_i_*, *kgen_i_*, and *km_i_* were kept constant during the global sensitivity analysis to gain a better understanding, since these parameters significantly influenced the outputs but do not contribute to improved insight into the biological mechanisms. From the analysis, it appeared that *CISH* KO has a central role in the production of IL-2, TNF-α, CTLA-4, PD1 and FasL, and it may interact with other parts of the pathway. An important observation is the influence of GSK3β in the expression of these proteins, indicating a possible additional target for multiplex gene-editing or drug targeting. GSK3β has been previously identified as a promising target for improving T cell activation, function and proliferation, despite the decrease in T cell migration[42], [43], [44], [82]. Furthermore, the steady-state simulations showed that CISH is the most crucial protein to target, but PD1 and GSK3β can also potentially be targeted, in conjunction with CISH targeting/knockout, to enhance TIL therapy effectiveness as well.

In conclusion, our model predicts that *CISH* KO is effective in increasing both activation and inhibitory markers. To compensate for the increase in inhibitory gene expression, a multiplex gene-editing approach could be explored, where *CISH* is knocked-out in conjunction with the knockout of *CTLA4*, *PDCD1*, *FAS*, and *GSK3B* to enhance T cell cytotoxicity and proliferation. While multiplex editing of *CISH*, *PDCD1*, *CTLA4*, *FAS*, and *GSK3B* may enhance TIL anti-tumor activity, simultaneous removal of multiple immune checkpoints and apoptotic controls can cause severe side effects such as uncontrolled T cell activation, autoimmunity lymphoproliferation[90], [91], [92], [93]. The two most significant limitations of the current model are the limited experimental data that were used to calibrate the model and the focus on NFAT. A next step for improving the current model would be to collect more experimental time series data and include the NFκB transcription factor.

## ACKNOWLEDGMENTS

B.R.W. acknowledges funding from Office of Discovery and Translation, NIH grants R21CA237789, R21AI163731, P01CA254849, P50CA136393, U54CA268069, R01AI146009, and Children’s Cancer Research Fund. B.S.M. acknowledges funding from NIH grants R01AI146009, R01AI161017, P01CA254849, P50CA136393, U24OD026641, U54CA232561, P30CA077598, U54CA268069, Children’s Cancer Research Fund, the Fanconi Anemia Research Fund, and the Randy Shaver Cancer and Community Fund. D.J.O. acknowledges funding support from NIH grants U54CA210190, U54CA268069, and P01CA254849.

## Supplementary Information

To estimate the baseline generation rates for mRNAs, we assumed that the average mRNA length is 1000 nucleotides, the maximum elongation rate 70 nt/s and the average number of RNA polymerases[89], [90], [91] in a T-cell is 300. Moreover, it was assumed that the diameter of a T-cell is 10 *μm*, thus its volume is equal to 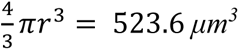[92].

**Figure A.1.**
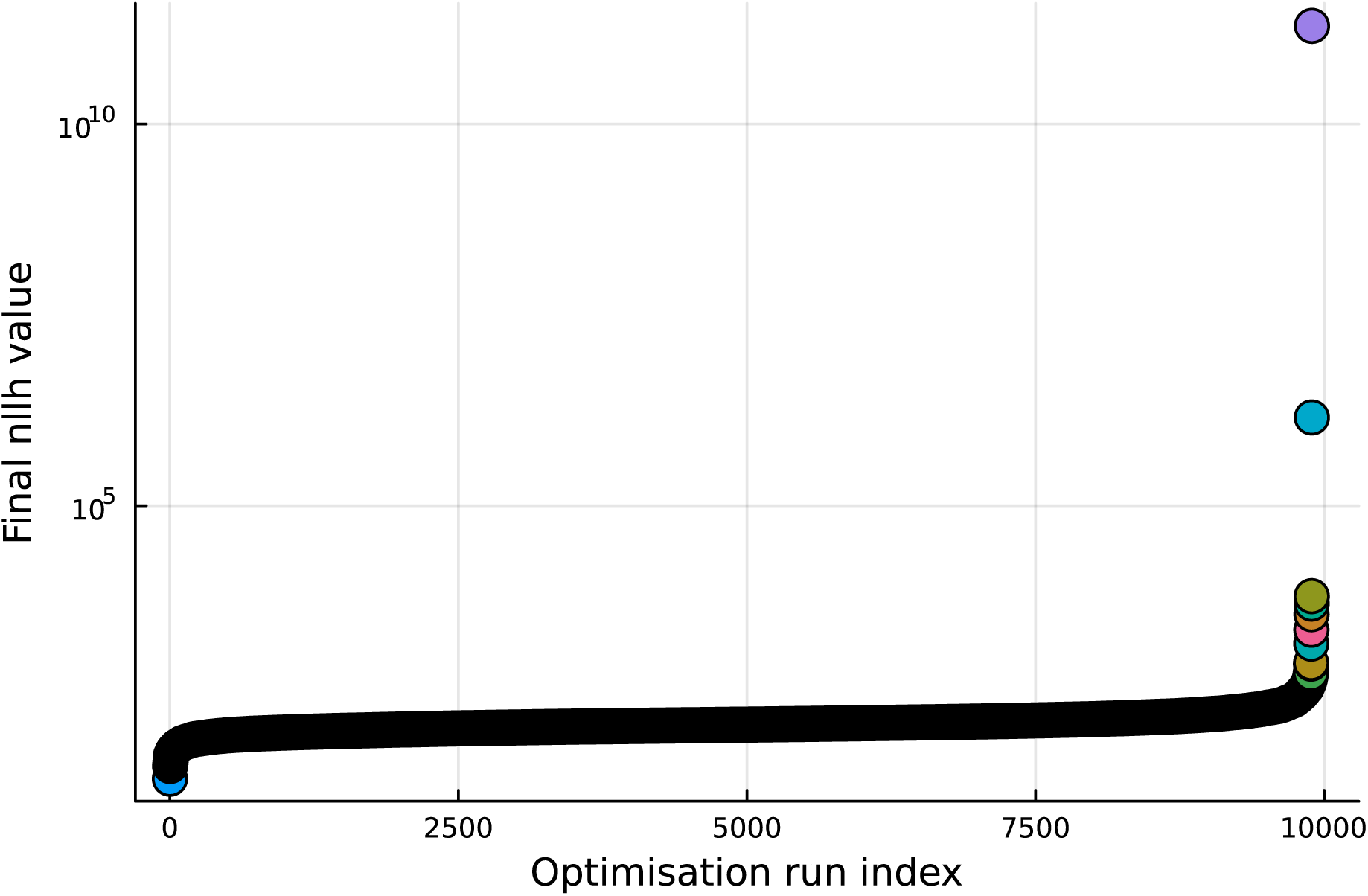
Waterfall plot for 10000 parameter estimations with different initial parameter values. The ranking is from the best on the left to the worst on the right on the x-axis. The y-axis corresponds to the negative log-likelihood, calculated based on (1).

**Table A.1.**
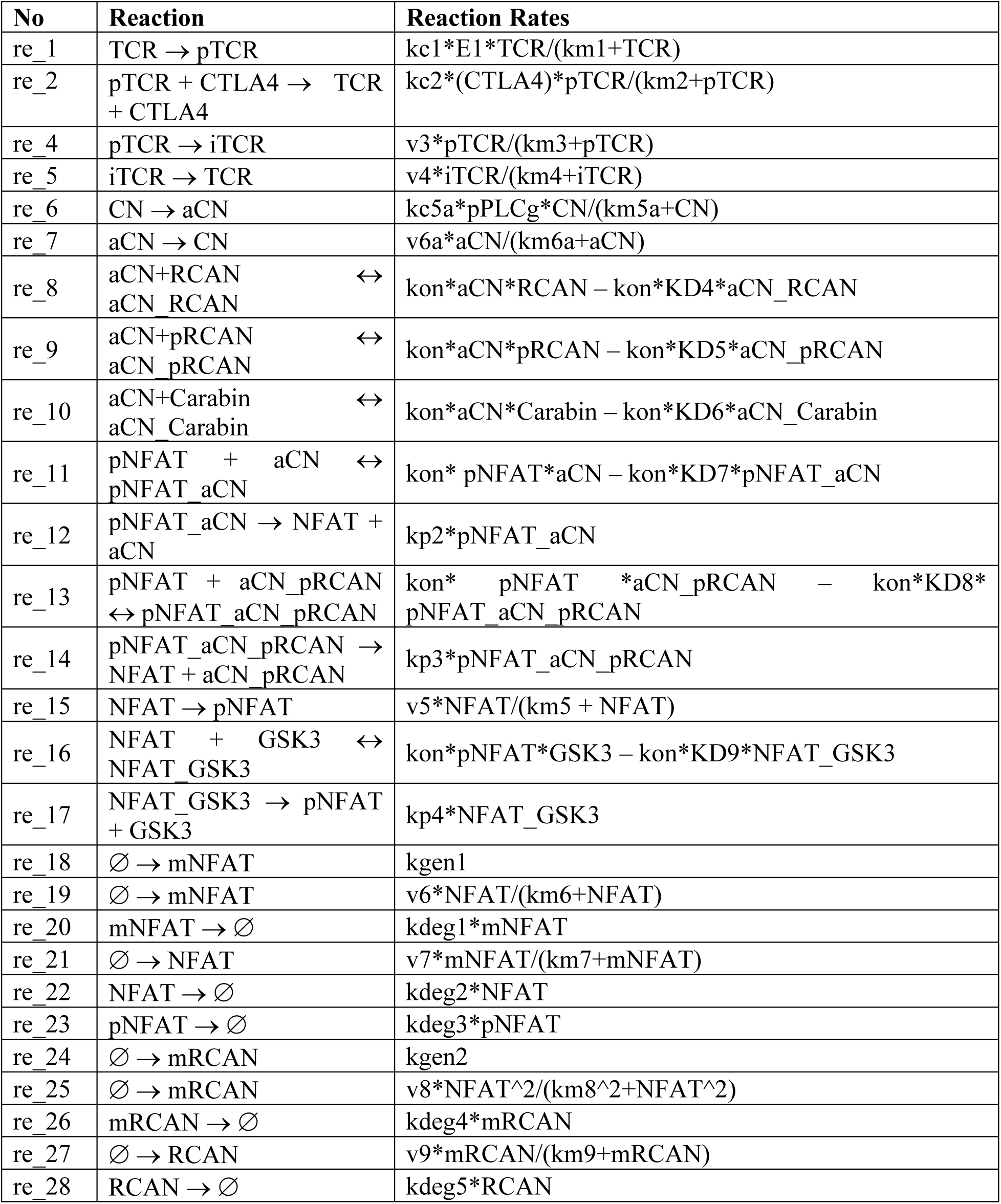
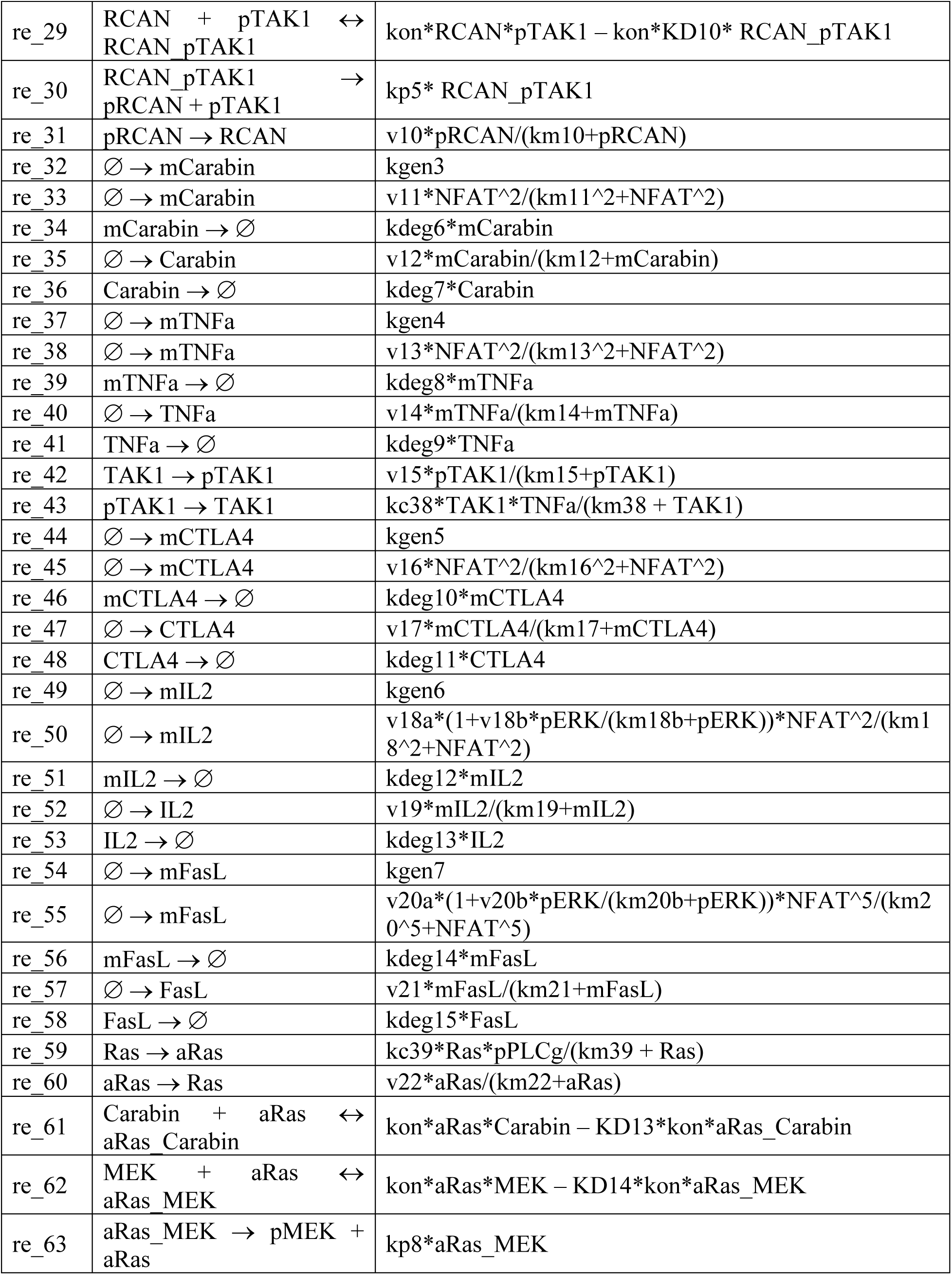
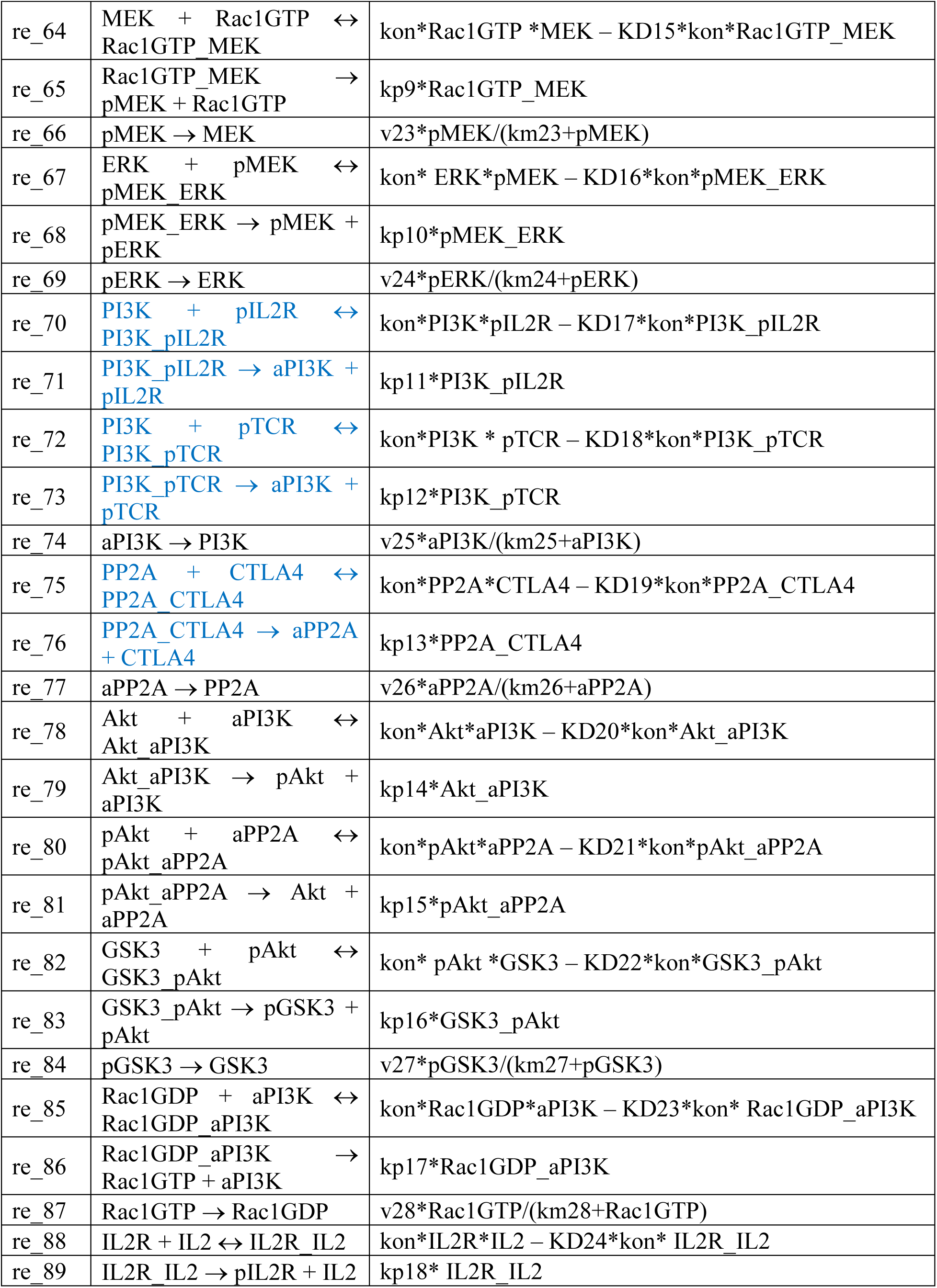
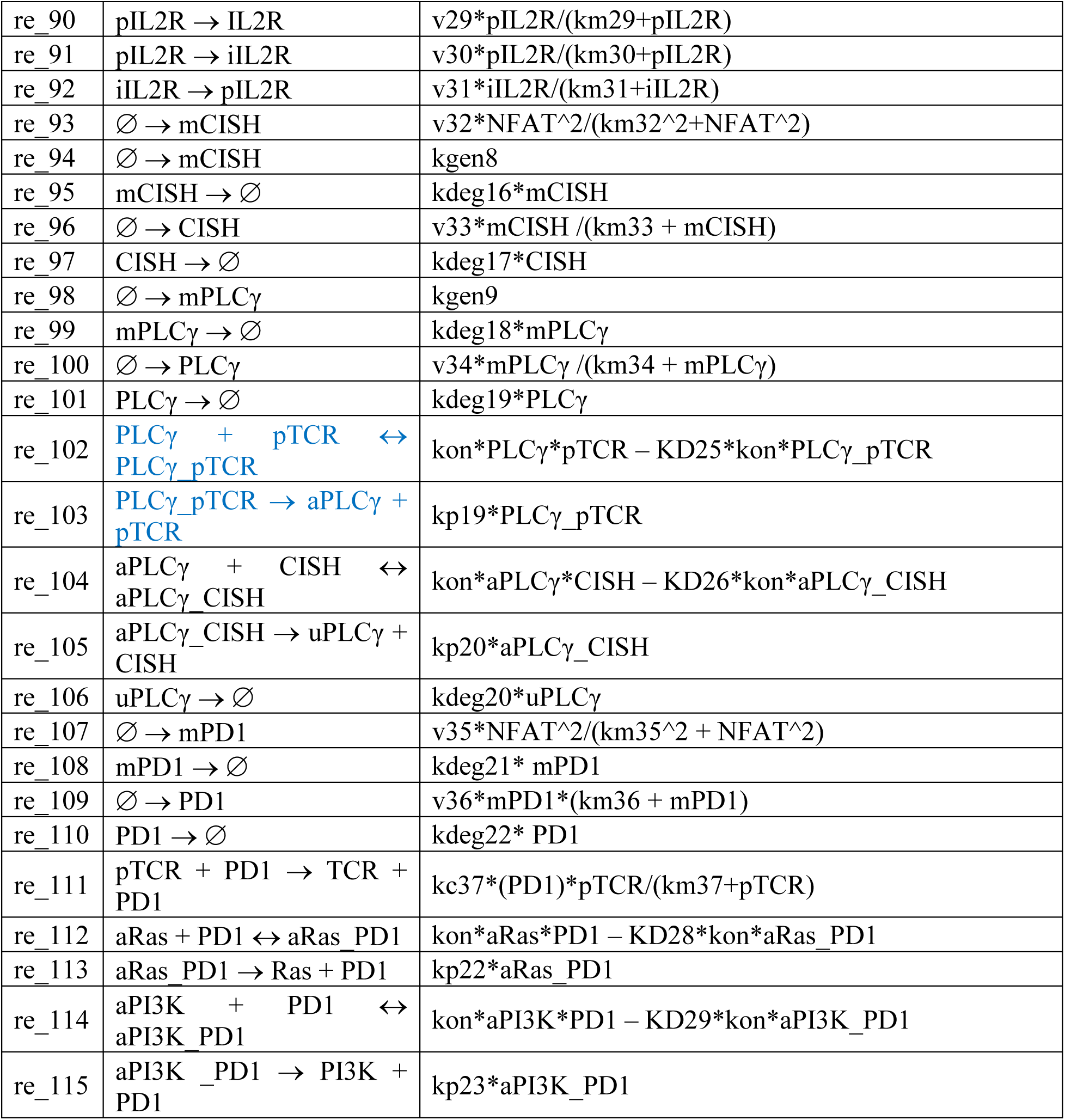
Table of reaction rates for CISH model.

It is assumed that the receptors TCR, CTLA-4, PD1 and IL2R have different binding motifs for different proteins, so the reactions with blue color do not affect the mass balance for the receptors.

**Table A.2.**
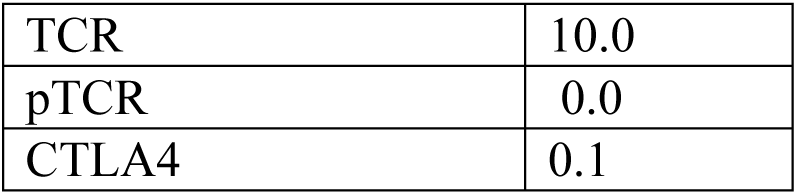
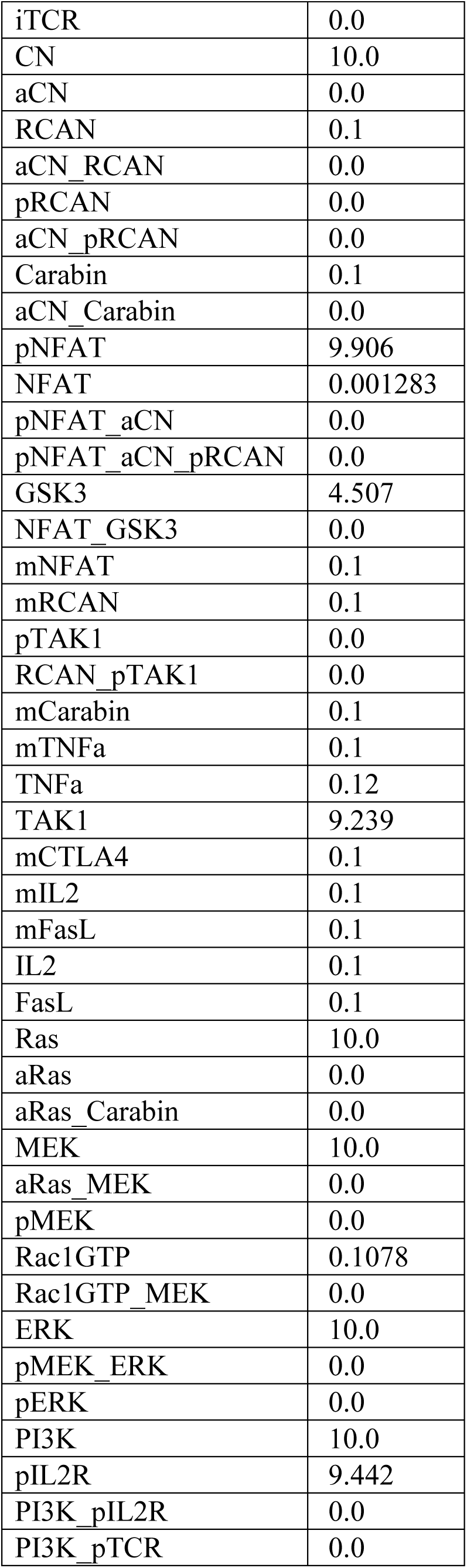
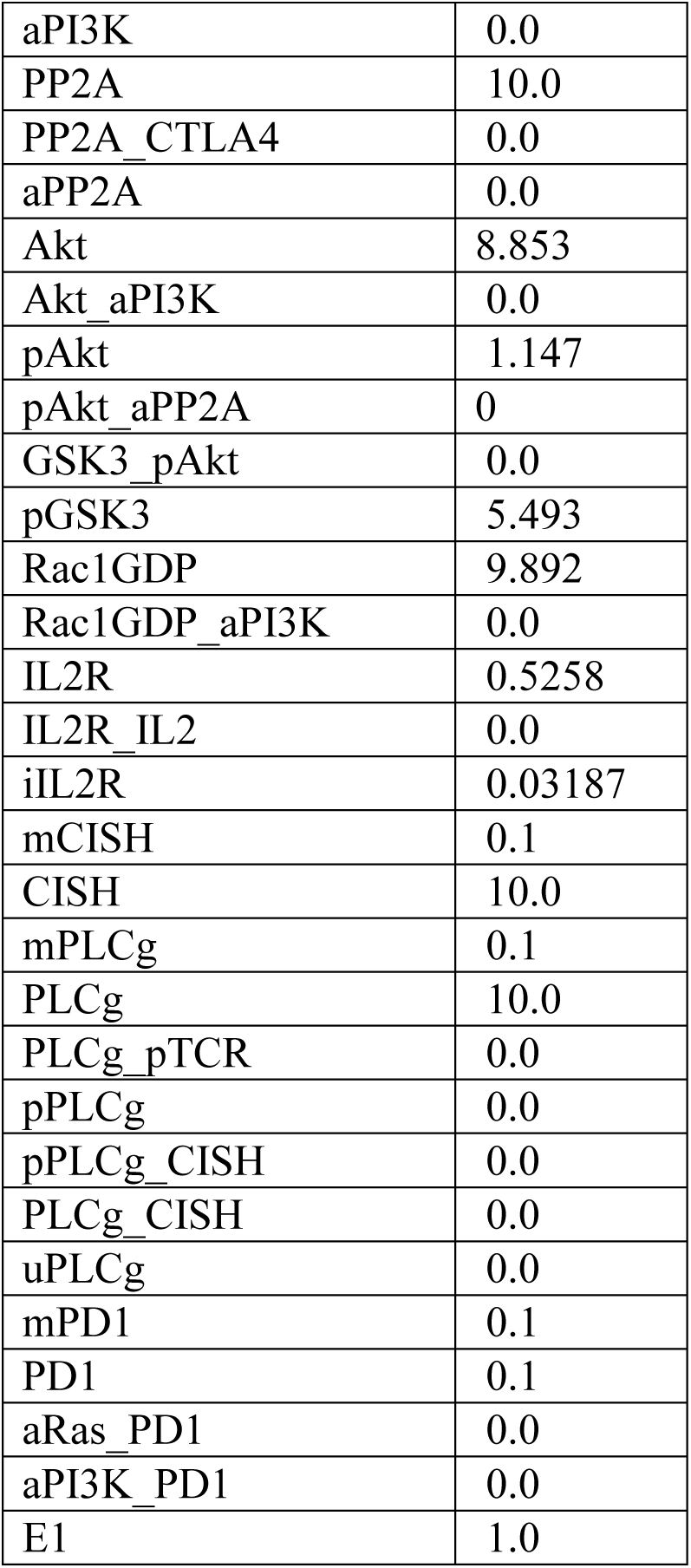
Table of the initial conditions (nM) for each species.

**Table A.3.**
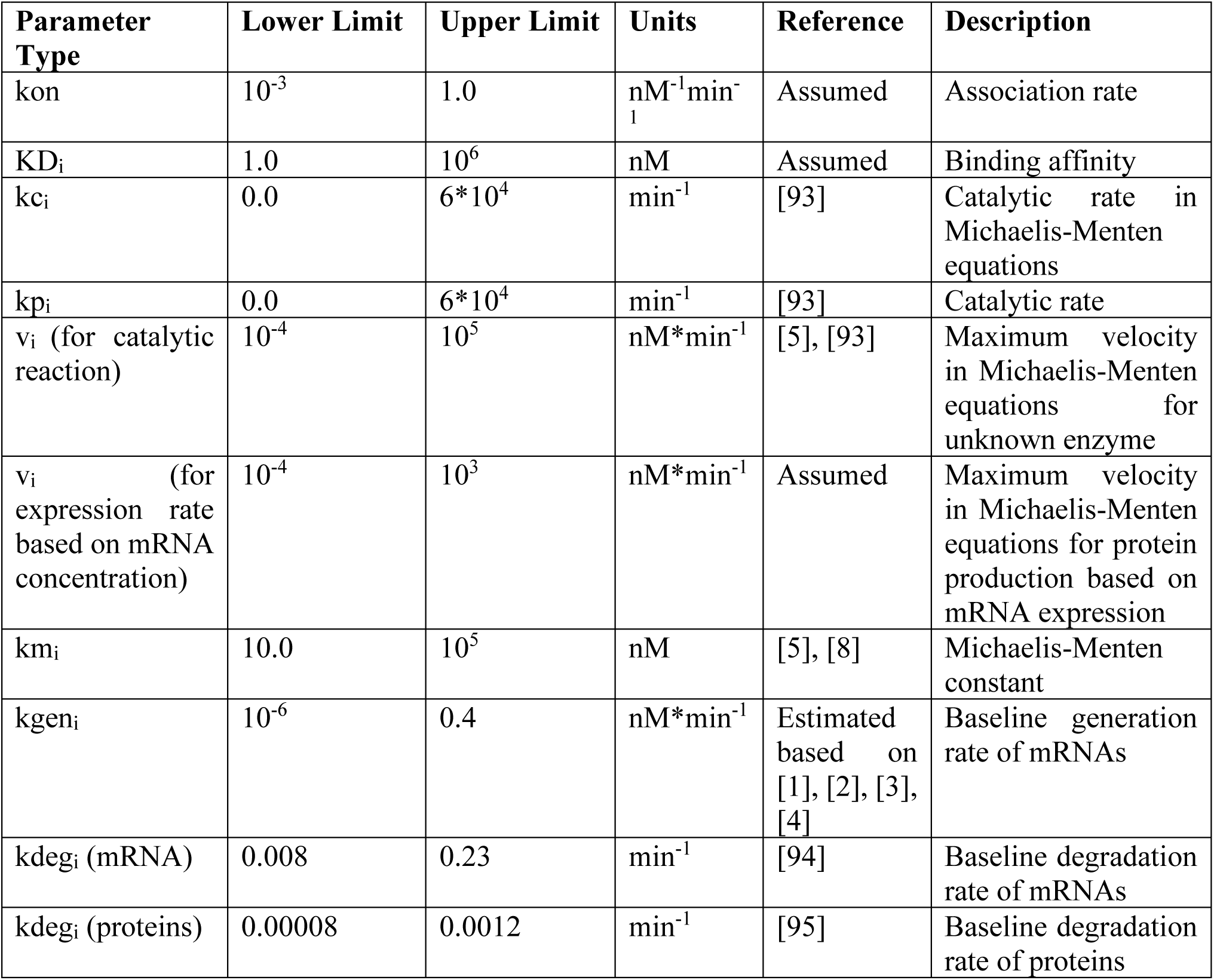
Upper and lower bounds of model parameters.

**Table A.4.**
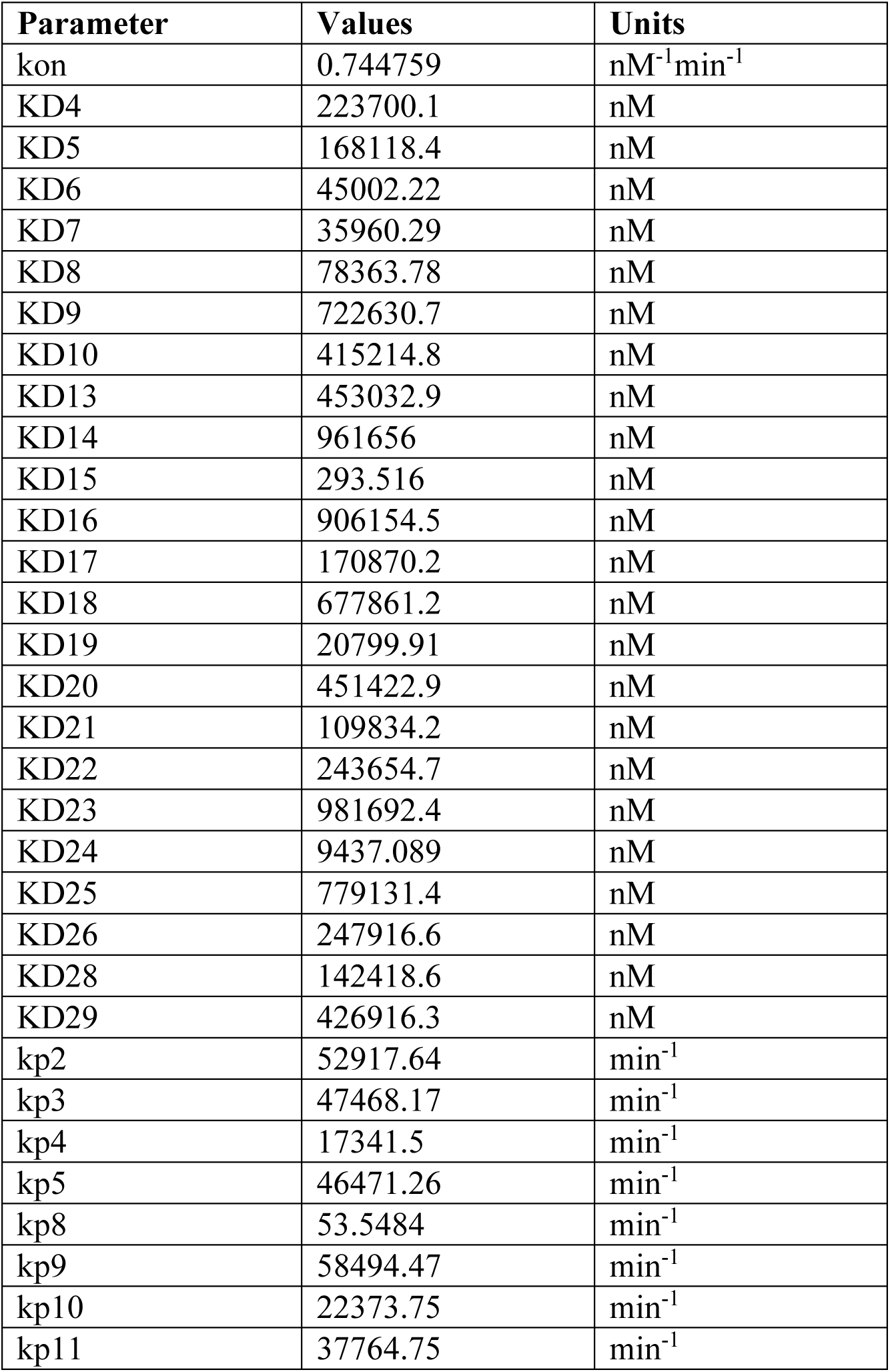
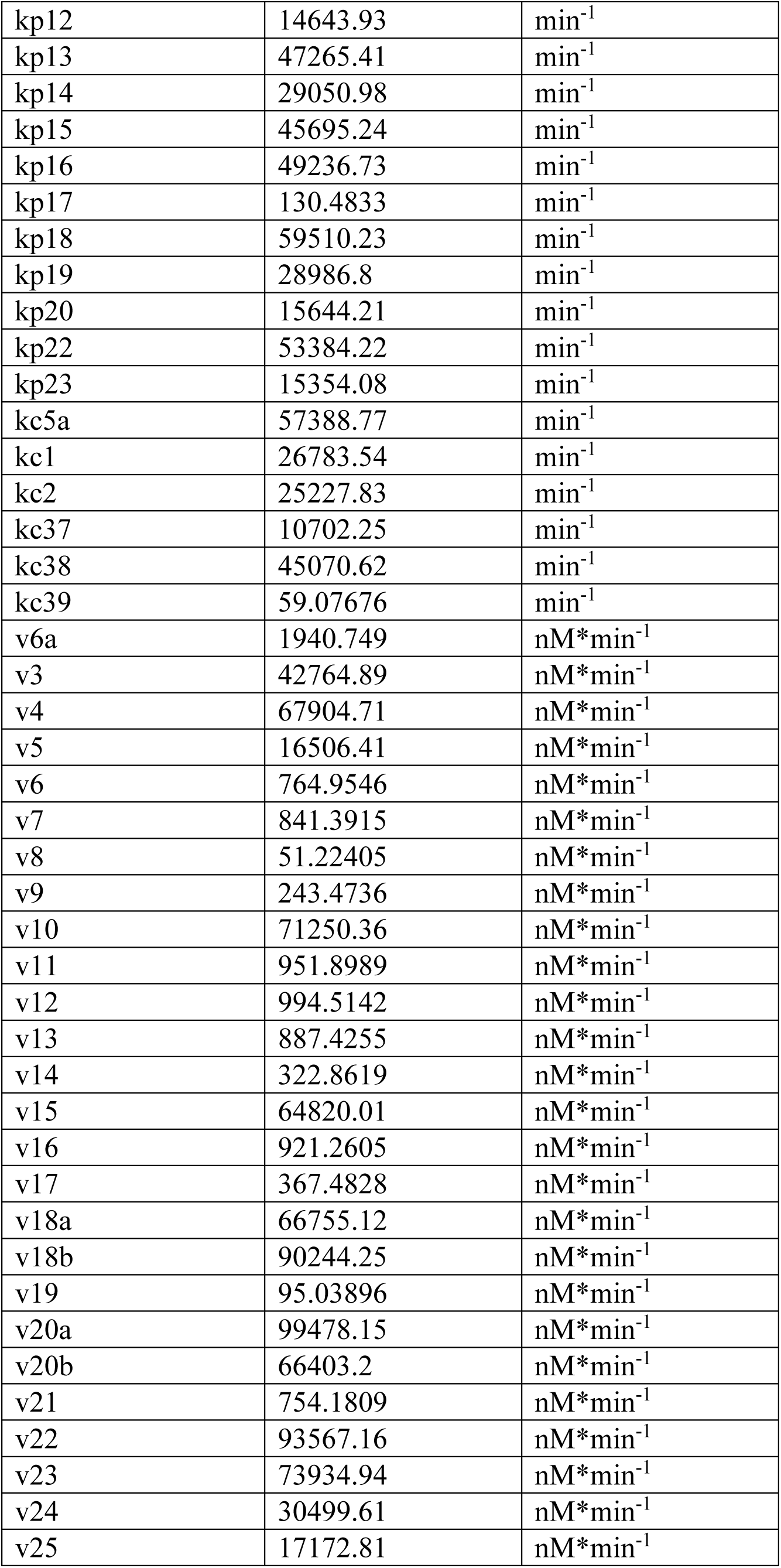
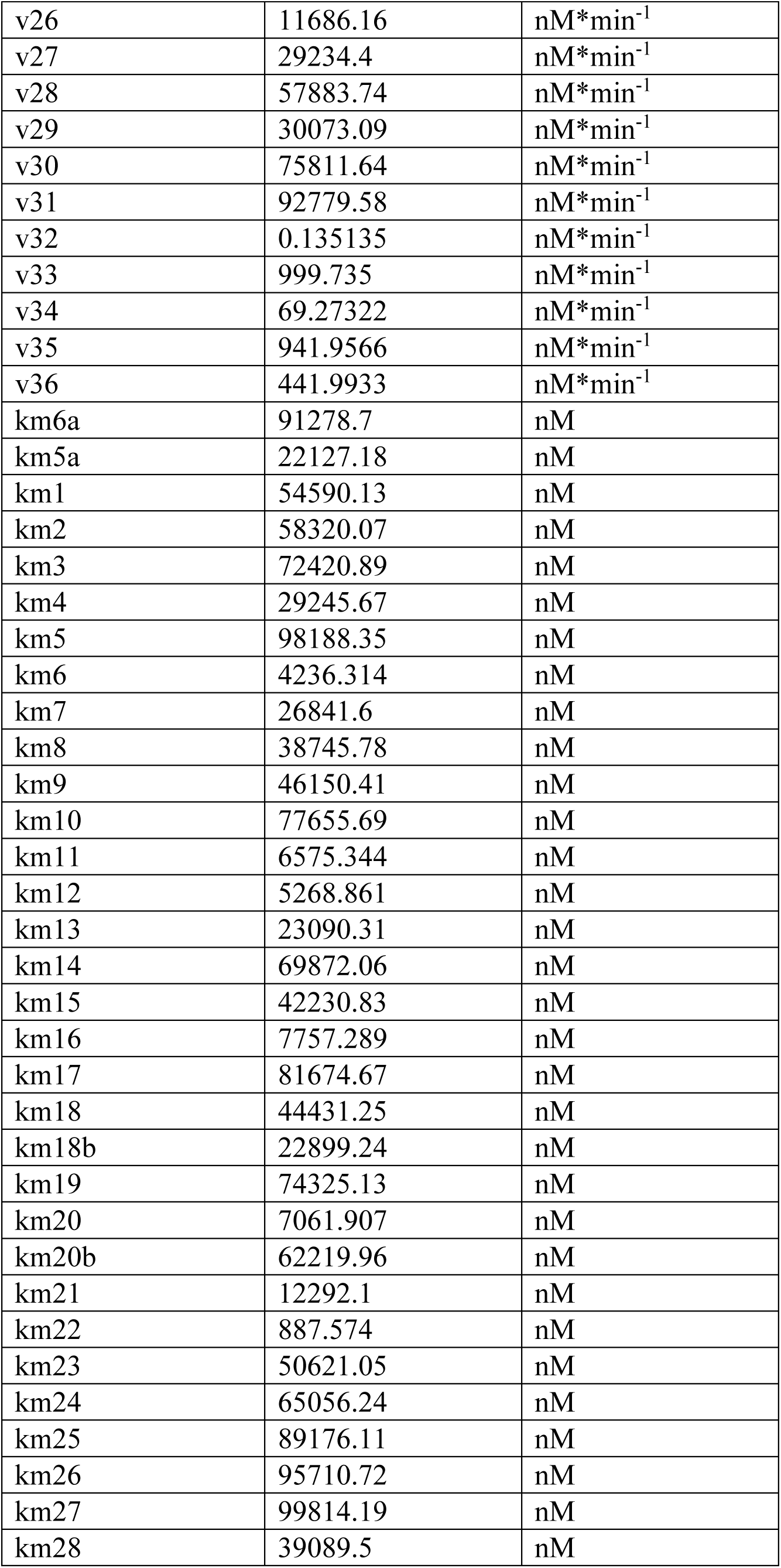
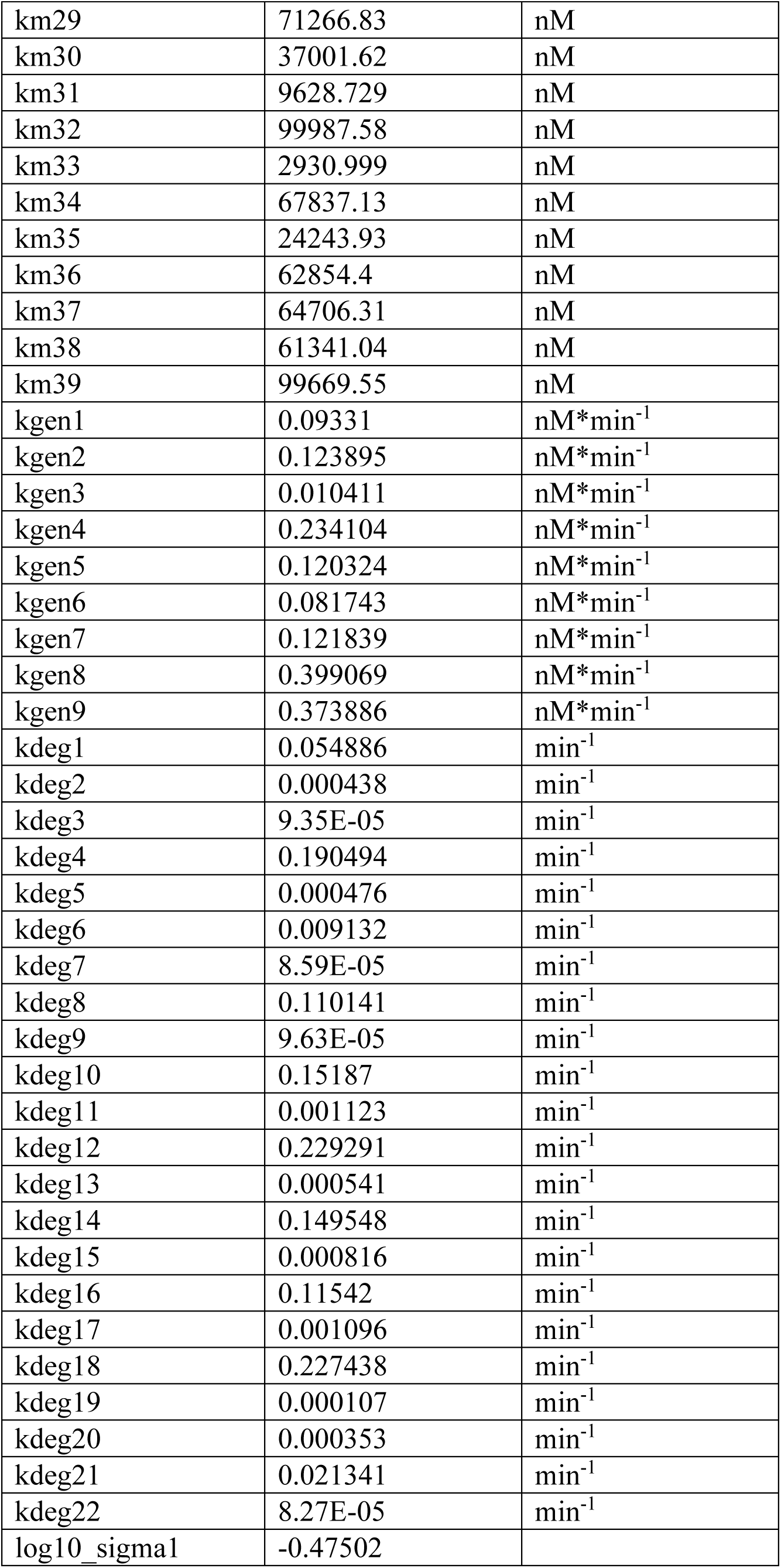
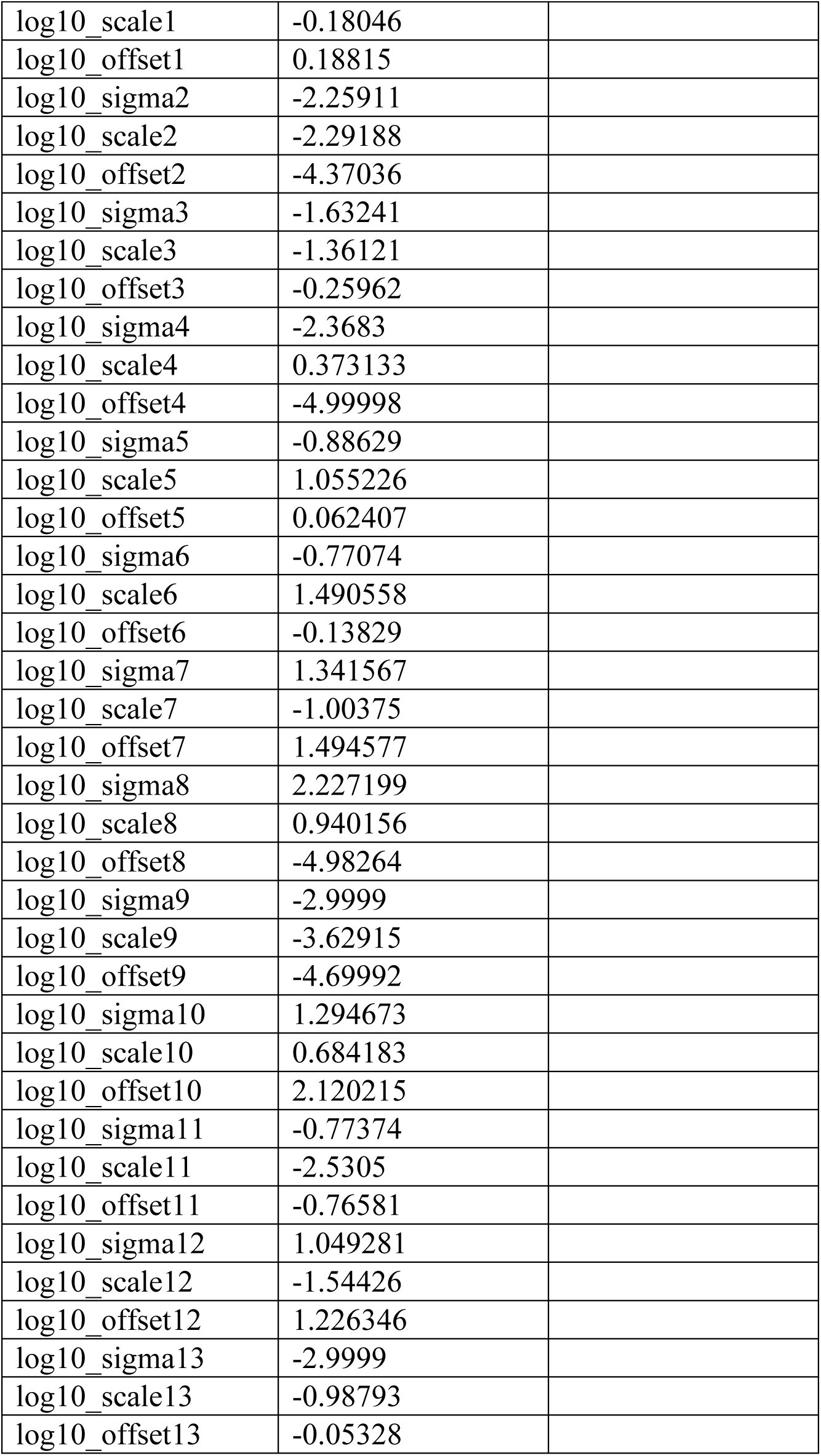
Fitted parameters.

