## Supplementary Information for "Mechanistic modeling predicts efficacy of CISH knockout in tumor-infiltrating lymphocytes with synergistic gene editing"

To estimate the baseline generation rates for mRNAs, we assumed that the average mRNA length is 1000 nucleotides, the maximum elongation rate 70 nt/s and the average number of RNA polymerases[89], [90], [91] in a T-cell is 300. Moreover, it was assumed that the diameter of a T-cell is 10 $\mu m$, thus its volume is equal to $\frac{4}{3}\pi r^3 = 523.6\ \mu m^3$[92].

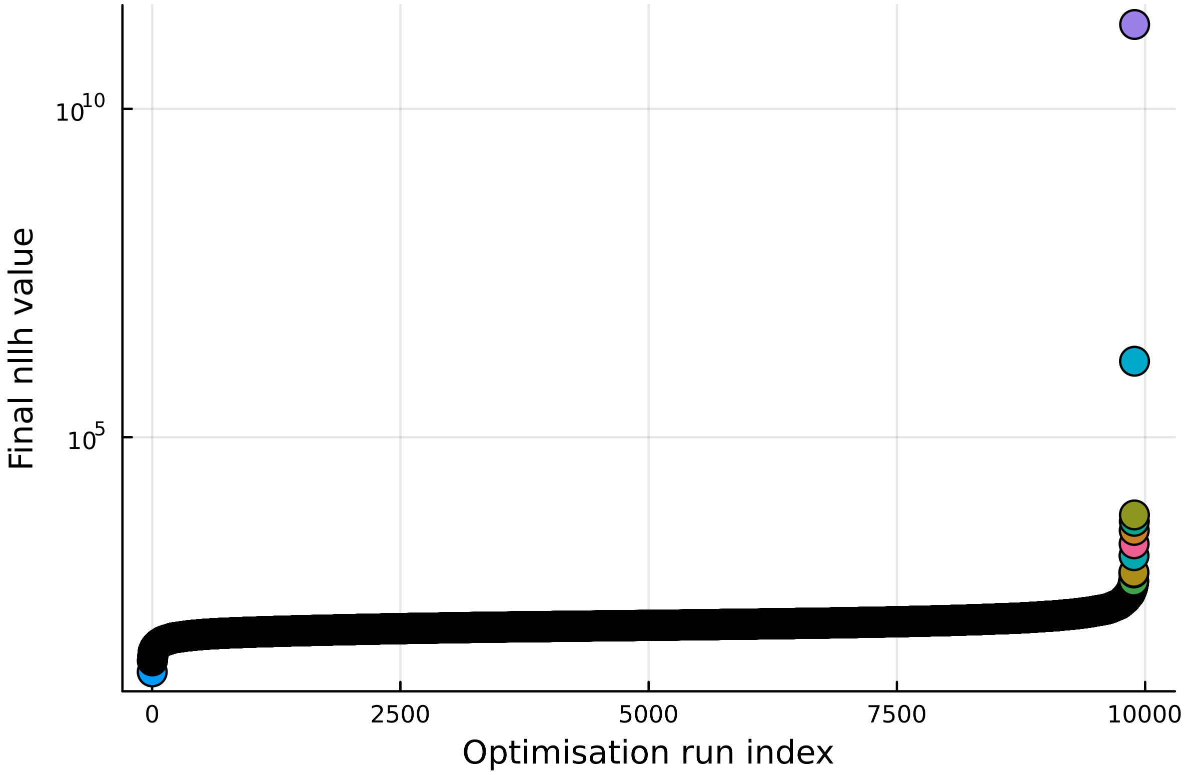

**Figure A. 1.** Waterfall plot for 10000 parameter estimations with different initial parameter values. The ranking is from the best on the left to the worst on the right on the x-axis. The y-axis corresponds to the negative log-likelihood, calculated based on (1).

**Table A. 1.** Table of reaction rates for CISH model.

| No | Reaction | Reaction Rates |
|---|---|---|
| re_1 | TCR → pTCR | kc1*E1*TCR/(km1+TCR) |
| re_2 | pTCR + CTLA4 → TCR + CTLA4 | kc2*(CTLA4)*pTCR/(km2+pTCR) |
| re_4 | pTCR → iTCR | v3*pTCR/(km3+pTCR) |
| re_5 | iTCR → TCR | v4*iTCR/(km4+iTCR) |
| re_6 | CN → aCN | kc5a*pPLCg*CN/(km5a+CN) |
| re_7 | aCN → CN | v6a*aCN/(km6a+aCN) |
| re_8 | aCN+RCAN ↔ aCN_RCAN | kon*aCN*RCAN – kon*KD4*aCN_RCAN |
| re_9 | aCN+pRCAN ↔ aCN_pRCAN | kon*aCN*pRCAN – kon*KD5*aCN_pRCAN |
| re_10 | aCN+Carabin ↔ aCN_Carabin | kon*aCN*Carabin – kon*KD6*aCN_Carabin |
| re_11 | pNFAT + aCN ↔ pNFAT_aCN | kon* pNFAT*aCN – kon*KD7*pNFAT_aCN |
| re_12 | pNFAT_aCN → NFAT + aCN | kp2*pNFAT_aCN |
| re_13 | pNFAT + aCN_pRCAN ↔ pNFAT_aCN_pRCAN | kon* pNFAT *aCN_pRCAN – kon*KD8* pNFAT_aCN_pRCAN |
| re_14 | pNFAT_aCN_pRCAN → NFAT + aCN_pRCAN | kp3*pNFAT_aCN_pRCAN |
| re_15 | NFAT → pNFAT | v5*NFAT/(km5 + NFAT) |
| re_16 | NFAT + GSK3 ↔ NFAT_GSK3 | kon*pNFAT*GSK3 – kon*KD9*NFAT_GSK3 |
| re_17 | NFAT_GSK3 → pNFAT + GSK3 | kp4*NFAT_GSK3 |
| re_18 | ∅ → mNFAT | kgen1 |
| re_19 | ∅ → mNFAT | v6*NFAT/(km6+NFAT) |
| re_20 | mNFAT → ∅ | kdeg1*mNFAT |
| re_21 | ∅ → NFAT | v7*mNFAT/(km7+mNFAT) |
| re_22 | NFAT → ∅ | kdeg2*NFAT |
| re_23 | pNFAT → ∅ | kdeg3*pNFAT |
| re_24 | ∅ → mRCAN | kgen2 |
| re_25 | ∅ → mRCAN | v8*NFAT^2/(km8^2+NFAT^2) |
| re_26 | mRCAN → ∅ | kdeg4*mRCAN |
| re_27 | ∅ → RCAN | v9*mRCAN/(km9+mRCAN) |
| re_28 | RCAN → ∅ | kdeg5*RCAN |

| | | |
|---|---|---|
| re_29 | RCAN + pTAK1 ↔ RCAN_pTAK1 | kon*RCAN*pTAK1 – kon*KD10* RCAN_pTAK1 |
| re_30 | RCAN_pTAK1 → pRCAN + pTAK1 | kp5* RCAN_pTAK1 |
| re_31 | pRCAN → RCAN | v10*pRCAN/(km10+pRCAN) |
| re_32 | ∅ → mCarabin | kgen3 |
| re_33 | ∅ → mCarabin | v11*NFAT^2/(km11^2+NFAT^2) |
| re_34 | mCarabin → ∅ | kdeg6*mCarabin |
| re_35 | ∅ → Carabin | v12*mCarabin/(km12+mCarabin) |
| re_36 | Carabin → ∅ | kdeg7*Carabin |
| re_37 | ∅ → mTNFa | kgen4 |
| re_38 | ∅ → mTNFa | v13*NFAT^2/(km13^2+NFAT^2) |
| re_39 | mTNFa → ∅ | kdeg8*mTNFa |
| re_40 | ∅ → TNFa | v14*mTNFa/(km14+mTNFa) |
| re_41 | TNFa → ∅ | kdeg9*TNFa |
| re_42 | TAK1 → pTAK1 | v15*pTAK1/(km15+pTAK1) |
| re_43 | pTAK1 → TAK1 | kc38*TAK1*TNFa/(km38 + TAK1) |
| re_44 | ∅ → mCTLA4 | kgen5 |
| re_45 | ∅ → mCTLA4 | v16*NFAT^2/(km16^2+NFAT^2) |
| re_46 | mCTLA4 → ∅ | kdeg10*mCTLA4 |
| re_47 | ∅ → CTLA4 | v17*mCTLA4/(km17+mCTLA4) |
| re_48 | CTLA4 → ∅ | kdeg11*CTLA4 |
| re_49 | ∅ → mIL2 | kgen6 |
| re_50 | ∅ → mIL2 | v18a*(1+v18b*pERK/(km18b+pERK))*NFAT^2/(km18^2+NFAT^2) |
| re_51 | mIL2 → ∅ | kdeg12*mIL2 |
| re_52 | ∅ → IL2 | v19*mIL2/(km19+mIL2) |
| re_53 | IL2 → ∅ | kdeg13*IL2 |
| re_54 | ∅ → mFasL | kgen7 |
| re_55 | ∅ → mFasL | v20a*(1+v20b*pERK/(km20b+pERK))*NFAT^5/(km20^5+NFAT^5) |
| re_56 | mFasL → ∅ | kdeg14*mFasL |
| re_57 | ∅ → FasL | v21*mFasL/(km21+mFasL) |
| re_58 | FasL → ∅ | kdeg15*FasL |
| re_59 | Ras → aRas | kc39*Ras*pPLCg/(km39 + Ras) |
| re_60 | aRas → Ras | v22*aRas/(km22+aRas) |
| re_61 | Carabin + aRas ↔ aRas_Carabin | kon*aRas*Carabin – KD13*kon*aRas_Carabin |
| re_62 | MEK + aRas ↔ aRas_MEK | kon*aRas*MEK – KD14*kon*aRas_MEK |
| re_63 | aRas_MEK → pMEK + aRas | kp8*aRas_MEK |

| | | |
|---|---|---|
| re_64 | MEK + Rac1GTP ↔ Rac1GTP_MEK | kon*Rac1GTP *MEK – KD15*kon*Rac1GTP_MEK |
| re_65 | Rac1GTP_MEK → pMEK + Rac1GTP | kp9*Rac1GTP_MEK |
| re_66 | pMEK → MEK | v23*pMEK/(km23+pMEK) |
| re_67 | ERK + pMEK ↔ pMEK_ERK | kon* ERK*pMEK – KD16*kon*pMEK_ERK |
| re_68 | pMEK_ERK → pMEK + pERK | kp10*pMEK_ERK |
| re_69 | pERK → ERK | v24*pERK/(km24+pERK) |
| re_70 | PI3K + pIL2R ↔ PI3K_pIL2R | kon*PI3K*pIL2R – KD17*kon*PI3K_pIL2R |
| re_71 | PI3K_pIL2R → aPI3K + pIL2R | kp11*PI3K_pIL2R |
| re_72 | PI3K + pTCR ↔ PI3K_pTCR | kon*PI3K * pTCR – KD18*kon*PI3K_pTCR |
| re_73 | PI3K_pTCR → aPI3K + pTCR | kp12*PI3K_pTCR |
| re_74 | aPI3K → PI3K | v25*aPI3K/(km25+aPI3K) |
| re_75 | PP2A + CTLA4 ↔ PP2A_CTLA4 | kon*PP2A*CTLA4 – KD19*kon*PP2A_CTLA4 |
| re_76 | PP2A_CTLA4 → aPP2A + CTLA4 | kp13*PP2A_CTLA4 |
| re_77 | aPP2A → PP2A | v26*aPP2A/(km26+aPP2A) |
| re_78 | Akt + aPI3K ↔ Akt_aPI3K | kon*Akt*aPI3K – KD20*kon*Akt_aPI3K |
| re_79 | Akt_aPI3K → pAkt + aPI3K | kp14*Akt_aPI3K |
| re_80 | pAkt + aPP2A ↔ pAkt_aPP2A | kon*pAkt*aPP2A – KD21*kon*pAkt_aPP2A |
| re_81 | pAkt_aPP2A → Akt + aPP2A | kp15*pAkt_aPP2A |
| re_82 | GSK3 + pAkt ↔ GSK3_pAkt | kon* pAkt *GSK3 – KD22*kon*GSK3_pAkt |
| re_83 | GSK3_pAkt → pGSK3 + pAkt | kp16*GSK3_pAkt |
| re_84 | pGSK3 → GSK3 | v27*pGSK3/(km27+pGSK3) |
| re_85 | Rac1GDP + aPI3K ↔ Rac1GDP_aPI3K | kon*Rac1GDP*aPI3K – KD23*kon* Rac1GDP_aPI3K |
| re_86 | Rac1GDP_aPI3K → Rac1GTP + aPI3K | kp17*Rac1GDP_aPI3K |
| re_87 | Rac1GTP → Rac1GDP | v28*Rac1GTP/(km28+Rac1GTP) |
| re_88 | IL2R + IL2 ↔ IL2R_IL2 | kon*IL2R*IL2 – KD24*kon* IL2R_IL2 |
| re_89 | IL2R_IL2 → pIL2R + IL2 | kp18* IL2R_IL2 |

| | | |
|---|---|---|
| re_90 | pIL2R → IL2R | v29*pIL2R/(km29+pIL2R) |
| re_91 | pIL2R → iIL2R | v30*pIL2R/(km30+pIL2R) |
| re_92 | iIL2R → pIL2R | v31*iIL2R/(km31+iIL2R) |
| re_93 | ∅ → mCISH | v32*NFAT^2/(km32^2+NFAT^2) |
| re_94 | ∅ → mCISH | kgen8 |
| re_95 | mCISH → ∅ | kdeg16*mCISH |
| re_96 | ∅ → CISH | v33*mCISH /(km33 + mCISH) |
| re_97 | CISH → ∅ | kdeg17*CISH |
| re_98 | ∅ → mPLCγ | kgen9 |
| re_99 | mPLCγ → ∅ | kdeg18*mPLCγ |
| re_100 | ∅ → PLCγ | v34*mPLCγ /(km34 + mPLCγ) |
| re_101 | PLCγ → ∅ | kdeg19*PLCγ |
| re_102 | PLCγ + pTCR ↔ PLCγ_pTCR | kon*PLCγ*pTCR – KD25*kon*PLCγ_pTCR |
| re_103 | PLCγ_pTCR → aPLCγ + pTCR | kp19*PLCγ_pTCR |
| re_104 | aPLCγ + CISH ↔ aPLCγ_CISH | kon*aPLCγ*CISH – KD26*kon*aPLCγ_CISH |
| re_105 | aPLCγ_CISH → uPLCγ + CISH | kp20*aPLCγ_CISH |
| re_106 | uPLCγ → ∅ | kdeg20*uPLCγ |
| re_107 | ∅ → mPD1 | v35*NFAT^2/(km35^2 + NFAT^2) |
| re_108 | mPD1 → ∅ | kdeg21* mPD1 |
| re_109 | ∅ → PD1 | v36*mPD1*(km36 + mPD1) |
| re_110 | PD1 → ∅ | kdeg22* PD1 |
| re_111 | pTCR + PD1 → TCR + PD1 | kc37*(PD1)*pTCR/(km37+pTCR) |
| re_112 | aRas + PD1 ↔ aRas_PD1 | kon*aRas*PD1 – KD28*kon*aRas_PD1 |
| re_113 | aRas_PD1 → Ras + PD1 | kp22*aRas_PD1 |
| re_114 | aPI3K + PD1 ↔ aPI3K_PD1 | kon*aPI3K*PD1 – KD29*kon*aPI3K_PD1 |
| re_115 | aPI3K _PD1 → PI3K + PD1 | kp23*aPI3K_PD1 |

It is assumed that the receptors TCR, CTLA-4, PD1 and IL2R have different binding motifs for different proteins, so the reactions with blue color do not affect the mass balance for the receptors.

**Table A.2.** Table of the initial conditions (nM) for each species.

| | |
|---|---|
| TCR | 10.0 |
| pTCR | 0.0 |
| CTLA4 | 0.1 |

| | |
|---|---|
| iTCR | 0.0 |
| CN | 10.0 |
| aCN | 0.0 |
| RCAN | 0.1 |
| aCN_RCAN | 0.0 |
| pRCAN | 0.0 |
| aCN_pRCAN | 0.0 |
| Carabin | 0.1 |
| aCN_Carabin | 0.0 |
| pNFAT | 9.906 |
| NFAT | 0.001283 |
| pNFAT_aCN | 0.0 |
| pNFAT_aCN_pRCAN | 0.0 |
| GSK3 | 4.507 |
| NFAT_GSK3 | 0.0 |
| mNFAT | 0.1 |
| mRCAN | 0.1 |
| pTAK1 | 0.0 |
| RCAN_pTAK1 | 0.0 |
| mCarabin | 0.1 |
| mTNFa | 0.1 |
| TNFa | 0.12 |
| TAK1 | 9.239 |
| mCTLA4 | 0.1 |
| mIL2 | 0.1 |
| mFasL | 0.1 |
| IL2 | 0.1 |
| FasL | 0.1 |
| Ras | 10.0 |
| aRas | 0.0 |
| aRas_Carabin | 0.0 |
| MEK | 10.0 |
| aRas_MEK | 0.0 |
| pMEK | 0.0 |
| Rac1GTP | 0.1078 |
| Rac1GTP_MEK | 0.0 |
| ERK | 10.0 |
| pMEK_ERK | 0.0 |
| pERK | 0.0 |
| PI3K | 10.0 |
| pIL2R | 9.442 |
| PI3K_pIL2R | 0.0 |
| PI3K_pTCR | 0.0 |

| | |
|---|---|
| aPI3K | 0.0 |
| PP2A | 10.0 |
| PP2A_CTLA4 | 0.0 |
| aPP2A | 0.0 |
| Akt | 8.853 |
| Akt_aPI3K | 0.0 |
| pAkt | 1.147 |
| pAkt_aPP2A | 0 |
| GSK3_pAkt | 0.0 |
| pGSK3 | 5.493 |
| Rac1GDP | 9.892 |
| Rac1GDP_aPI3K | 0.0 |
| IL2R | 0.5258 |
| IL2R_IL2 | 0.0 |
| iIL2R | 0.03187 |
| mCISH | 0.1 |
| CISH | 10.0 |
| mPLCg | 0.1 |
| PLCg | 10.0 |
| PLCg_pTCR | 0.0 |
| pPLCg | 0.0 |
| pPLCg_CISH | 0.0 |
| PLCg_CISH | 0.0 |
| uPLCg | 0.0 |
| mPD1 | 0.1 |
| PD1 | 0.1 |
| aRas_PD1 | 0.0 |
| aPI3K_PD1 | 0.0 |
| E1 | 1.0 |

**Table A.3.** Upper and lower bounds of model parameters.

| Parameter Type | Lower Limit | Upper Limit | Units | Reference | Description |
|---|---|---|---|---|---|
| kon | $10^{-3}$ | 1.0 | $nM^{-1}min^{-1}$ | Assumed | Association rate |
| $KD_i$ | 1.0 | $10^6$ | nM | Assumed | Binding affinity |
| $kc_i$ | 0.0 | $6*10^4$ | $min^{-1}$ | [93] | Catalytic rate in Michaelis-Menten equations |
| $kp_i$ | 0.0 | $6*10^4$ | $min^{-1}$ | [93] | Catalytic rate |
| $v_i$ (for catalytic reaction) | $10^{-4}$ | $10^5$ | $nM*min^{-1}$ | [5], [93] | Maximum velocity in Michaelis-Menten equations for unknown enzyme |
| $v_i$ (for expression rate based on mRNA concentration) | $10^{-4}$ | $10^3$ | $nM*min^{-1}$ | Assumed | Maximum velocity in Michaelis-Menten equations for protein production based on mRNA expression |
| $km_i$ | 10.0 | $10^5$ | nM | [5], [8] | Michaelis-Menten constant |
| $kgen_i$ | $10^{-6}$ | 0.4 | $nM*min^{-1}$ | Estimated based on [1], [2], [3], [4] | Baseline generation rate of mRNAs |
| $kdeg_i$ (mRNA) | 0.008 | 0.23 | $min^{-1}$ | [94] | Baseline degradation rate of mRNAs |
| $kdeg_i$ (proteins) | 0.00008 | 0.0012 | $min^{-1}$ | [95] | Baseline degradation rate of proteins |

**Table A.4.** Fitted parameters.

| Parameter | Values | Units |
|---|---|---|
| kon | 0.744759 | $nM^{-1}min^{-1}$ |
| KD4 | 223700.1 | nM |
| KD5 | 168118.4 | nM |
| KD6 | 45002.22 | nM |
| KD7 | 35960.29 | nM |
| KD8 | 78363.78 | nM |
| KD9 | 722630.7 | nM |
| KD10 | 415214.8 | nM |
| KD13 | 453032.9 | nM |
| KD14 | 961656 | nM |
| KD15 | 293.516 | nM |
| KD16 | 906154.5 | nM |
| KD17 | 170870.2 | nM |
| KD18 | 677861.2 | nM |
| KD19 | 20799.91 | nM |
| KD20 | 451422.9 | nM |
| KD21 | 109834.2 | nM |
| KD22 | 243654.7 | nM |
| KD23 | 981692.4 | nM |
| KD24 | 9437.089 | nM |
| KD25 | 779131.4 | nM |
| KD26 | 247916.6 | nM |
| KD28 | 142418.6 | nM |
| KD29 | 426916.3 | nM |
| kp2 | 52917.64 | $min^{-1}$ |
| kp3 | 47468.17 | $min^{-1}$ |
| kp4 | 17341.5 | $min^{-1}$ |
| kp5 | 46471.26 | $min^{-1}$ |
| kp8 | 53.5484 | $min^{-1}$ |
| kp9 | 58494.47 | $min^{-1}$ |
| kp10 | 22373.75 | $min^{-1}$ |
| kp11 | 37764.75 | $min^{-1}$ |

| | | |
|---|---|---|
| kp12 | 14643.93 | $min^{-1}$ |
| kp13 | 47265.41 | $min^{-1}$ |
| kp14 | 29050.98 | $min^{-1}$ |
| kp15 | 45695.24 | $min^{-1}$ |
| kp16 | 49236.73 | $min^{-1}$ |
| kp17 | 130.4833 | $min^{-1}$ |
| kp18 | 59510.23 | $min^{-1}$ |
| kp19 | 28986.8 | $min^{-1}$ |
| kp20 | 15644.21 | $min^{-1}$ |
| kp22 | 53384.22 | $min^{-1}$ |
| kp23 | 15354.08 | $min^{-1}$ |
| kc5a | 57388.77 | $min^{-1}$ |
| kc1 | 26783.54 | $min^{-1}$ |
| kc2 | 25227.83 | $min^{-1}$ |
| kc37 | 10702.25 | $min^{-1}$ |
| kc38 | 45070.62 | $min^{-1}$ |
| kc39 | 59.07676 | $min^{-1}$ |
| v6a | 1940.749 | $nM*min^{-1}$ |
| v3 | 42764.89 | $nM*min^{-1}$ |
| v4 | 67904.71 | $nM*min^{-1}$ |
| v5 | 16506.41 | $nM*min^{-1}$ |
| v6 | 764.9546 | $nM*min^{-1}$ |
| v7 | 841.3915 | $nM*min^{-1}$ |
| v8 | 51.22405 | $nM*min^{-1}$ |
| v9 | 243.4736 | $nM*min^{-1}$ |
| v10 | 71250.36 | $nM*min^{-1}$ |
| v11 | 951.8989 | $nM*min^{-1}$ |
| v12 | 994.5142 | $nM*min^{-1}$ |
| v13 | 887.4255 | $nM*min^{-1}$ |
| v14 | 322.8619 | $nM*min^{-1}$ |
| v15 | 64820.01 | $nM*min^{-1}$ |
| v16 | 921.2605 | $nM*min^{-1}$ |
| v17 | 367.4828 | $nM*min^{-1}$ |
| v18a | 66755.12 | $nM*min^{-1}$ |
| v18b | 90244.25 | $nM*min^{-1}$ |
| v19 | 95.03896 | $nM*min^{-1}$ |
| v20a | 99478.15 | $nM*min^{-1}$ |
| v20b | 66403.2 | $nM*min^{-1}$ |
| v21 | 754.1809 | $nM*min^{-1}$ |
| v22 | 93567.16 | $nM*min^{-1}$ |
| v23 | 73934.94 | $nM*min^{-1}$ |
| v24 | 30499.61 | $nM*min^{-1}$ |
| v25 | 17172.81 | $nM*min^{-1}$ |

| | | |
|---|---|---|
| v26 | 11686.16 | nM*min$^{-1}$ |
| v27 | 29234.4 | nM*min$^{-1}$ |
| v28 | 57883.74 | nM*min$^{-1}$ |
| v29 | 30073.09 | nM*min$^{-1}$ |
| v30 | 75811.64 | nM*min$^{-1}$ |
| v31 | 92779.58 | nM*min$^{-1}$ |
| v32 | 0.135135 | nM*min$^{-1}$ |
| v33 | 999.735 | nM*min$^{-1}$ |
| v34 | 69.27322 | nM*min$^{-1}$ |
| v35 | 941.9566 | nM*min$^{-1}$ |
| v36 | 441.9933 | nM*min$^{-1}$ |
| km6a | 91278.7 | nM |
| km5a | 22127.18 | nM |
| km1 | 54590.13 | nM |
| km2 | 58320.07 | nM |
| km3 | 72420.89 | nM |
| km4 | 29245.67 | nM |
| km5 | 98188.35 | nM |
| km6 | 4236.314 | nM |
| km7 | 26841.6 | nM |
| km8 | 38745.78 | nM |
| km9 | 46150.41 | nM |
| km10 | 77655.69 | nM |
| km11 | 6575.344 | nM |
| km12 | 5268.861 | nM |
| km13 | 23090.31 | nM |
| km14 | 69872.06 | nM |
| km15 | 42230.83 | nM |
| km16 | 7757.289 | nM |
| km17 | 81674.67 | nM |
| km18 | 44431.25 | nM |
| km18b | 22899.24 | nM |
| km19 | 74325.13 | nM |
| km20 | 7061.907 | nM |
| km20b | 62219.96 | nM |
| km21 | 12292.1 | nM |
| km22 | 887.574 | nM |
| km23 | 50621.05 | nM |
| km24 | 65056.24 | nM |
| km25 | 89176.11 | nM |
| km26 | 95710.72 | nM |
| km27 | 99814.19 | nM |
| km28 | 39089.5 | nM |

| | | |
|---|---|---|
| km29 | 71266.83 | nM |
| km30 | 37001.62 | nM |
| km31 | 9628.729 | nM |
| km32 | 99987.58 | nM |
| km33 | 2930.999 | nM |
| km34 | 67837.13 | nM |
| km35 | 24243.93 | nM |
| km36 | 62854.4 | nM |
| km37 | 64706.31 | nM |
| km38 | 61341.04 | nM |
| km39 | 99669.55 | nM |
| kgen1 | 0.09331 | $nM*min^{-1}$ |
| kgen2 | 0.123895 | $nM*min^{-1}$ |
| kgen3 | 0.010411 | $nM*min^{-1}$ |
| kgen4 | 0.234104 | $nM*min^{-1}$ |
| kgen5 | 0.120324 | $nM*min^{-1}$ |
| kgen6 | 0.081743 | $nM*min^{-1}$ |
| kgen7 | 0.121839 | $nM*min^{-1}$ |
| kgen8 | 0.399069 | $nM*min^{-1}$ |
| kgen9 | 0.373886 | $nM*min^{-1}$ |
| kdeg1 | 0.054886 | $min^{-1}$ |
| kdeg2 | 0.000438 | $min^{-1}$ |
| kdeg3 | 9.35E-05 | $min^{-1}$ |
| kdeg4 | 0.190494 | $min^{-1}$ |
| kdeg5 | 0.000476 | $min^{-1}$ |
| kdeg6 | 0.009132 | $min^{-1}$ |
| kdeg7 | 8.59E-05 | $min^{-1}$ |
| kdeg8 | 0.110141 | $min^{-1}$ |
| kdeg9 | 9.63E-05 | $min^{-1}$ |
| kdeg10 | 0.15187 | $min^{-1}$ |
| kdeg11 | 0.001123 | $min^{-1}$ |
| kdeg12 | 0.229291 | $min^{-1}$ |
| kdeg13 | 0.000541 | $min^{-1}$ |
| kdeg14 | 0.149548 | $min^{-1}$ |
| kdeg15 | 0.000816 | $min^{-1}$ |
| kdeg16 | 0.11542 | $min^{-1}$ |
| kdeg17 | 0.001096 | $min^{-1}$ |
| kdeg18 | 0.227438 | $min^{-1}$ |
| kdeg19 | 0.000107 | $min^{-1}$ |
| kdeg20 | 0.000353 | $min^{-1}$ |
| kdeg21 | 0.021341 | $min^{-1}$ |
| kdeg22 | 8.27E-05 | $min^{-1}$ |
| log10_sigma1 | -0.47502 | |

| | |
|---|---|
| log10_scale1 | -0.18046 |
| log10_offset1 | 0.18815 |
| log10_sigma2 | -2.25911 |
| log10_scale2 | -2.29188 |
| log10_offset2 | -4.37036 |
| log10_sigma3 | -1.63241 |
| log10_scale3 | -1.36121 |
| log10_offset3 | -0.25962 |
| log10_sigma4 | -2.3683 |
| log10_scale4 | 0.373133 |
| log10_offset4 | -4.99998 |
| log10_sigma5 | -0.88629 |
| log10_scale5 | 1.055226 |
| log10_offset5 | 0.062407 |
| log10_sigma6 | -0.77074 |
| log10_scale6 | 1.490558 |
| log10_offset6 | -0.13829 |
| log10_sigma7 | 1.341567 |
| log10_scale7 | -1.00375 |
| log10_offset7 | 1.494577 |
| log10_sigma8 | 2.227199 |
| log10_scale8 | 0.940156 |
| log10_offset8 | -4.98264 |
| log10_sigma9 | -2.9999 |
| log10_scale9 | -3.62915 |
| log10_offset9 | -4.69992 |
| log10_sigma10 | 1.294673 |
| log10_scale10 | 0.684183 |
| log10_offset10 | 2.120215 |
| log10_sigma11 | -0.77374 |
| log10_scale11 | -2.5305 |
| log10_offset11 | -0.76581 |
| log10_sigma12 | 1.049281 |
| log10_scale12 | -1.54426 |
| log10_offset12 | 1.226346 |
| log10_sigma13 | -2.9999 |
| log10_scale13 | -0.98793 |
| log10_offset13 | -0.05328 |
